# Strong reduction of the chain rigidity of hyaluronan by selective binding of Ca^2+^ ions

**DOI:** 10.1101/2020.09.01.277194

**Authors:** G. Giubertoni, A. Pérez de Alba Ortíz, F. Bano, X. Zhang, R.J. Linhardt, D. E. Green, P. L. DeAngelis, G.H. Koenderink, R. P. Richter, B. Ensing, H.J. Bakker

## Abstract

The biological functions of natural polyelectrolytes are strongly influenced by the presence of ions, which bind to the polymer chains and thereby modify their properties. Although the biological impact of such modifications is well-recognized, a detailed molecular picture of the binding process and of the mechanisms that drive the subsequent structural changes in the polymer is lacking. Here, we study the molecular mechanism of the condensation of calcium, a divalent cation, on hyaluronan, a ubiquitous polymer in human tissues. By combining two-dimensional infrared spectroscopy experiments with molecular dynamics simulations, we find that calcium specifically binds to hyaluronan at millimolar concentrations. Because of its large size and charge, the calcium cation can bind simultaneously to the negatively charged carboxylate group and the amide group of adjacent saccharide units. Molecular dynamics simulations and single-chain force spectroscopy measurements provide evidence that the binding of the calcium ions weakens the intra-molecular hydrogen-bond network of hyaluronan, increasing the flexibility of the polymer chain. We also observe that the binding of calcium to hyaluronan saturates at a maximum binding fraction of ~10-15 mol %. This saturation indicates that the binding of Ca^2+^ strongly reduces the probability of subsequent binding of Ca^2+^ at neighboring binding sites, possibly as a result of enhanced conformational fluctuations and/or electrostatic repulsion effects. Our findings provide a detailed molecular picture of ion condensation, and reveal the severe effect of a few, selective and localized electrostatic interactions on the rigidity of a polyelectrolyte chain.

**TOC:** 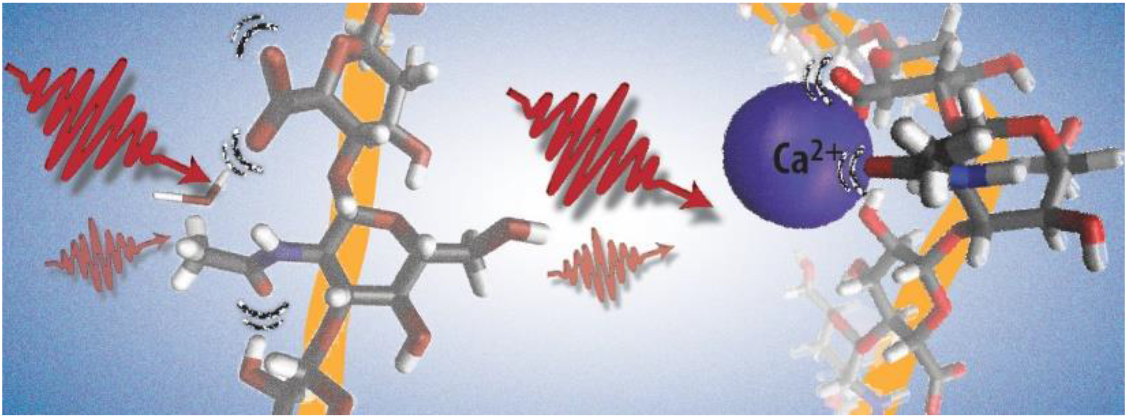

---

Polyelectrolytes are charged polymers, which are widely present in nature and in man-made materials for applications ranging from wound dressing to oil-recovery.^1,2^ Because of their charged nature, the conformation and physical properties of polyelectrolyte chains strongly depend on the solution pH and on the salt conditions. The electrostatic repulsive forces among the charges along the chain enhance the polymer rigidity. Usually, dissolved ions are mobile, and can thus screen the charges along the chain, thereby reducing the total persistence length that is a measure of the chain rigidity.^3^ Localized electrostatic interactions between ions and the charges on the chain (normally referred to as Manning condensation) can occur if the distance between the charges on the chain is less than the Bjerrum length (λ_B_).^4^ One possible consequence is that the polymer backbone wraps around bound ions.^5,6^ Condensation of multivalent ions has also been reported to entail local ion “jackets”, and consequently, a reduction in persistence length of polyelectrolyte chains can also happen without creating local wrapping of the chain.^7^ Although the effect of condensation on the configuration of polyelectrolytes has been thoroughly studied in previous work,^5–8^ the molecular details of the complexes formed between cations and polyelectrolytes, and the molecular mechanisms underlying the conformational changes that follow from the ion binding, are still unknown.

Among all the natural polyelectrolytes, a specific class of extracellular matrix polysaccharides, the glycosaminoglycans (GAGs), is arguably one of the most important for animal life. In the human body, GAGs are critically important in many biological processes, such as proliferation, anticoagulation,^9,10^ inflammatory responses,^11,12^ and the immune response to external pathogens.^13^ Hyaluronan (HA) is the structurally simplest member of the GAG family. Alone or together with other extracellular matrix macromolecules (*e.g*., collagen), it dictates tissue elasticity, hydration and permeability, and it also directs cell behavior through multivalent engagement with cell surface receptors, such as CD44.^14^ These properties allow HA to mediate diverse functions in a wide range of physiological and pathological processes, including development,^15,16^ mammalian reproduction,^17^ inflammation,^18^ and tissue lubrication.^19,20^ Diseases such as cancer or osteoarthritis are correlated with changes in the average molecular weight, supramolecular organization and concentration of hyaluronan.^21–23^ Thanks to its biocompatibility, hyaluronan is also widely applied as a building block for responsive and biocompatible hydrogels.^24–26^

An interesting feature of hyaluronan is the sensitivity of its mechanical properties to a particular divalent cation, calcium. Calcium ions have been found to show an unusually strong effect on the thickness of hyaluronan brushes (*i.e.* dense arrays of hyaluronan chains grafted with one end to a surface) designed to emulate certain properties of the glycocalyx of cells.^27^ Moreover, at a concentration of a few mM of calcium ions, hyaluronan solutions show a decrease in viscosity that is much more drastic than in the presence of a similar concentration of sodium or magnesium.^28,29^ Consistent with this finding, a previous study also showed that calcium ion levels in the low mM range cause a reduced translational diffusivity of small solutes, such as glucose and lysine, in hyaluronan polymer solutions.^30^ These findings suggest that hyaluronan polymers change their molecular conformation when interacting with calcium ions.

Hyaluronan is a linear polysaccharide composed of repeating disaccharide units (Fig. 1a) made of *N*-acetylglucosamine (with an amide group), and glucuronic acid (with a carboxyl group) monosaccharides linked together by alternating β1 → 4 and β1 → 3 glycosidic linkages. The contour length per disaccharide is 1.0 nm. The distance between negative charges (at the carboxyl groups) is thus smaller than the Bjerrum length for divalent cations (~1.4 nm),^7,27^ and ion condensation may occur, leading to a reduction of the charge density along the chain.

**Figure 1:**
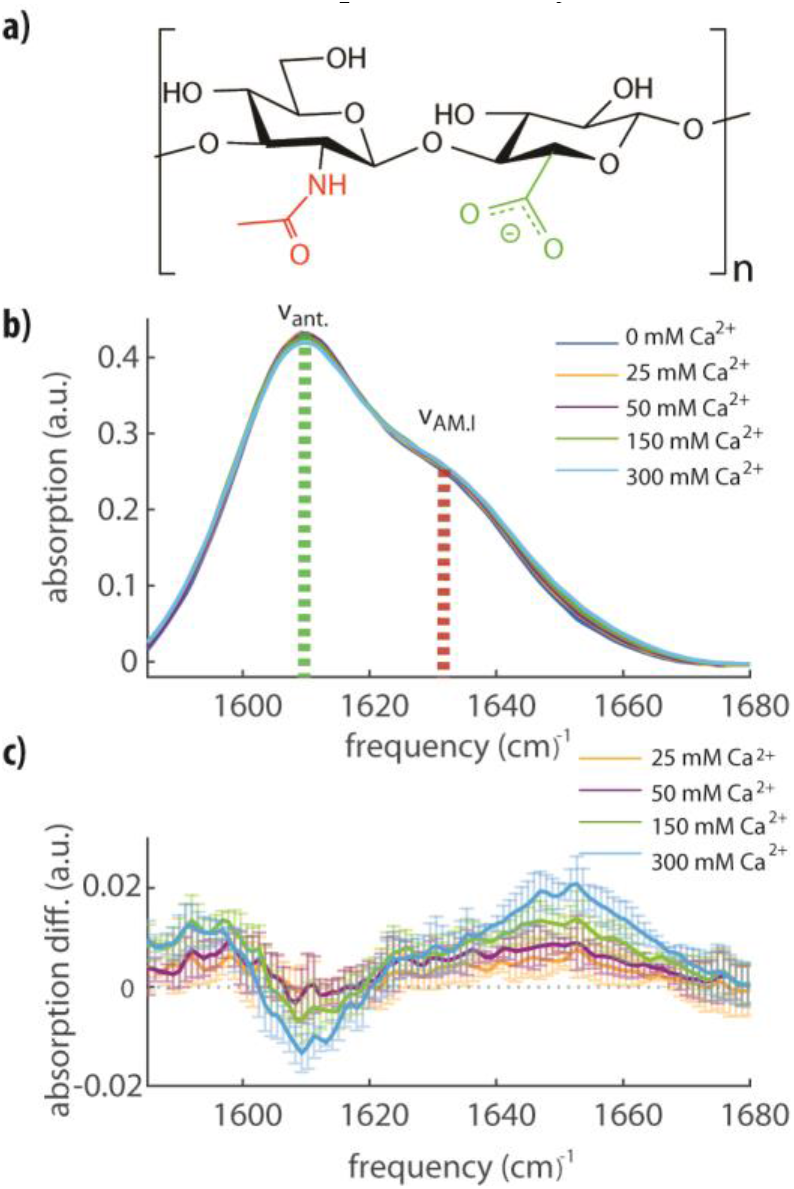
a) Molecular structure of a disaccharide unit of hyaluronan containing amide (*red*) and carboxylate (*green*) groups on adjacent saccharide units. b) Linear FTIR infrared spectra for a solution of hyaluronan at 20 mg/ml in water containing 0, 25, 50, 150 or 300 mM of calcium ions (as indicated). We observe the absorption peaks of the anti-symmetric stretching mode of the carboxylate anion group (ν_ant_.), and of the amide I vibration (ν_AM.I_). All the spectra are background subtracted. c) Differential FTIR infrared spectra obtained by subtracting the infrared spectrum in pure water (0 mM Ca^2+^) from the other spectra shown in b).

Here, we investigate the interaction between hyaluronan and calcium ions at the molecular level, with linear infrared (IR) spectroscopy, femtosecond two-dimensional infrared (2DIR) spectroscopy, single molecule force spectroscopy, and molecular dynamics simulations. We find that calcium ions bind to hyaluronan at millimolar concentrations, forming complexes with the carboxylate anion and amide groups. We further find that the formation of only a few of these complexes per any one polymer chain suffices to change the intramolecular hydrogen bond network and, as a result, the persistence length of hyaluronan polymers. We thus obtain a direct molecular picture of the binding mechanism between calcium and hyaluronan, and a molecular explanation of the change in flexibility of the polymer upon interaction with calcium ions.

## RESULTS

### Complexation of calcium ions with hyaluronan

We used linear and nonlinear IR spectroscopy to characterize the interaction between calcium ions and HA (see Supplementary Methods for details). In Fig. 1b we report linear infrared absorption spectra of a solution of HA at a concentration of 20 mg/ml, where we vary the CaCl_2_ concentration from 0 to 300 mM. In the frequency region between 1580 and 1680 cm^−1^, we observe two bands, one at 1609 cm^−1^ and the other at 1633 cm^−1^. Following the literature, we assign the band at 1609 cm^−1^ to the absorption band of the anti-symmetric stretching mode of the carboxylate anion group (ν_ant_), and the band at 1633 cm^−1^ to the absorption band of the amide I vibration (ν_AM.I_) of the amide group.^31^ Upon addition of CaCl_2_, we observe a small increase of the absorption around 1590 cm^−1^, and a decrease of the absorption near 1607 cm^−1^, indicating that the calcium affects the molecular vibrations of the carboxylate anion group. Most notable, however, is the enhanced absorption in the high-frequency region of the spectrum, corresponding to the high-frequency wing of the amide I band (1650 cm^−1^). In the absence of calcium, the amide group is hydrated on average by two water molecules, and each hydrogen bond induces a red-shift of the amide I absorption band of 10 to 20 cm^−1^.^32,33^ The observed partial blue shift of the amide I band from 1633 cm^−1^ to 1650 cm^−1^ thus suggests that calcium ions dehydrate a part of the amide groups. Computational studies have shown that in simple model systems containing a single amide group, calcium ions have a significant probability to be located close to the carbonyl oxygen (< 2.5 Å), and thus to be in direct contact with the amide group.^32–34^ The creation of such a cation-amide pair induces a red-shift in the amide I vibration with respect to the frequency of this mode in the gas phase, but this red-shift is smaller than the red-shift that results from the formation of hydrogen bonds with water molecules. In a previous study, calcium was found to bind to the carbonyl oxygen in a collinear fashion, displacing both water molecules,^32^ thus corroborating that the observed blue-shift of the amide vibration results from the binding of calcium ions.

At calcium concentrations below 150 mM, there is no significant difference between the linear absorption spectra measured with and without added salt. Nevertheless, based on previous reports, we do expect a significant effect of calcium ions on the hyaluronan structure already in this low-concentration regime. We studied the molecular-scale effect of calcium at low concentrations with two-dimensional infrared (2DIR) spectroscopy. 2DIR is a nonlinear technique in which spectroscopy the signal is linearly proportional to the vibrational cross-section (α ~ σ). Therefore, 2DIR is ideally suited to distinguish species with high cross-sections and low concentration (e.g., molecular vibrations of amide and carboxylate groups) from a background of species with low cross-sections and high concentrations (e.g., molecular vibrations of water). In Fig. 2a, we present 2DIR spectra measured for a hyaluronan solution with 0 and 25 mM calcium ions (Fig. S1 shows additional data spanning from 0 to 300 mM). In both spectra, we observe a strong signal when exciting at a pump frequency of 1607 cm^−1^, which extends to higher molecular vibrations are excited from the ground state (*n =* 0) to the first excited level (*n =* 1) with an intense femtosecond mid-infrared light pulse (pump pulse). This excitation leads to a change of the absorption of the excited and other vibrations that we probe with a second, weaker, broadband, femtosecond mid-infrared light pulse (probe pulse). The absorption change Δα measured with 2DIR is proportional to the square of the vibrational cross-section, σ (Δα ~ σ^2^), while in linear infrared probe frequencies and shows a shoulder at 1630 cm^−1^. The peak and the shoulder colored in blue represent a decrease in absorption due to bleaching of the fundamental *n* = 0 to *n* = 1 transitions of the ν_ant_ and ν_AM.I_ vibrations, respectively. The signals at lower probe frequencies colored in red represent the induced absorption of the *n* = 1 to *n* = 2 transition. Upon addition of calcium ions, we observe an enhanced absorption at higher frequencies (indicated by the arrow in Fig. 2b around 1660 cm^−1^), which can be seen more clearly in the lower part of Fig. 2b where we show the difference between the two slices taken along the diagonal of the bleach (also shown in Fig. S1), illustrating the enhanced absorption.

**Figure 2:**
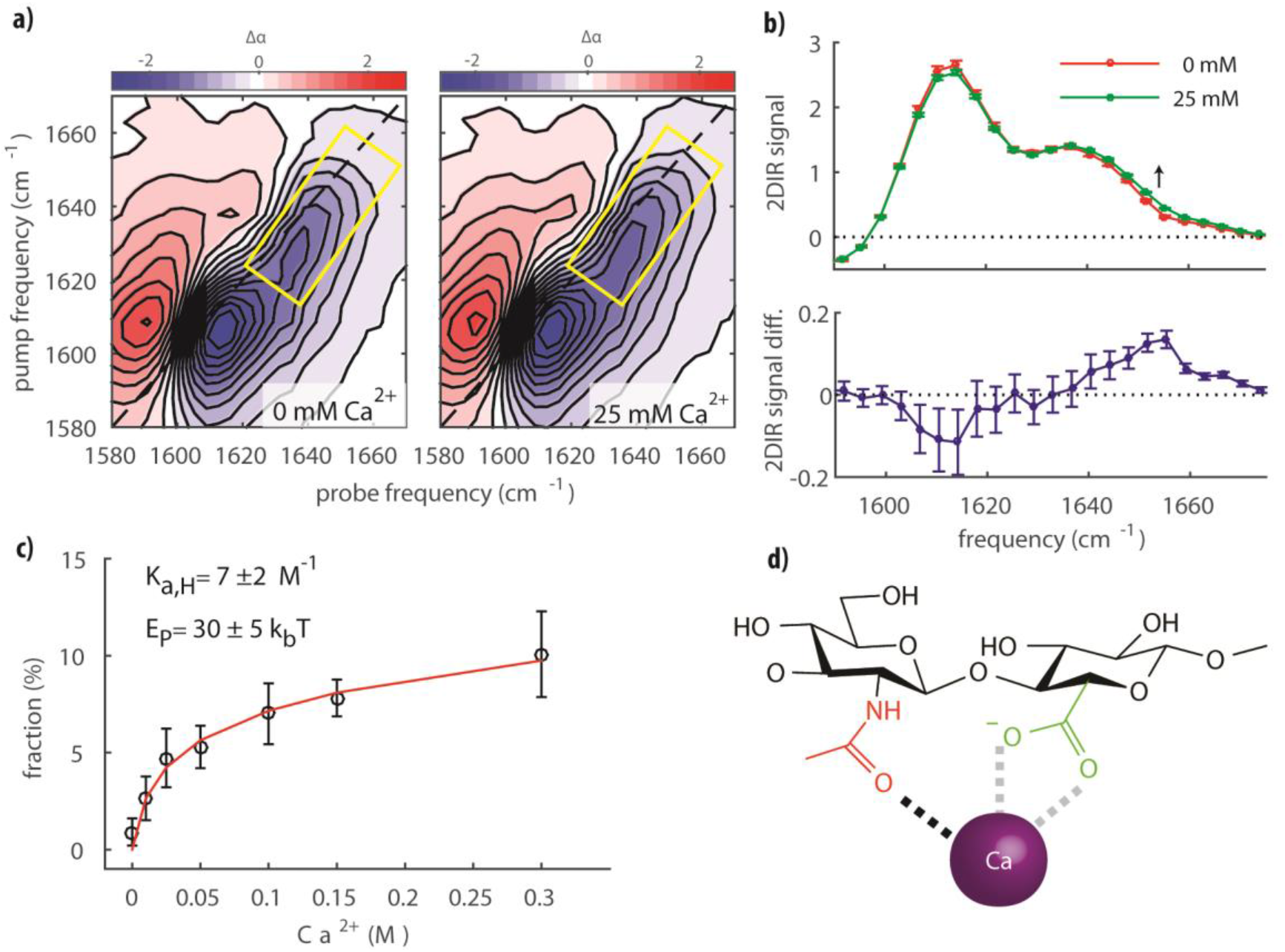
a) 2DIR spectra of hyaluronan at a concentration of 20 mg/ml in water containing 0 (*left*) and 25 mM (*right*) calcium ions. The waiting time between the pump and probe pulse was 0.3 ps. The *yellow rectangles* indicate the regions with the largest changes in absorption. b) Top: transient absorption 2DIR signals taken along the diagonal slice (*dashed line*: guideline for the eye set at 2DIR signal equal to 0) of the bleach in (a). Bottom: Differential spectrum obtained by subtracting the spectrum measured for a solution without calcium (0 mM) from the spectrum of a solution with 25 mM calcium ions. c) Fraction of complexed amide groups as a function of calcium concentration. The data (*symbols with error bars*) are fitted with a model that includes an increasing energy penalty with increasing occupation of the amide groups (see main text and Supplementary Methods for details). The mean values and error bars (which represent the standard deviation) were obtained by averaging over three different experiments. d) Illustration of the complex formed. The *grey dashed lines* show the bidentate binding of the carboxylate anion group to Ca^2+^.

We observe a significantly increased absorption on the blue side of the amide vibration peak already at a calcium concentration of 25 mM. As with the linear absorption spectra, we assign this enhanced absorption to the complexation of the amide carbonyl to the calcium ion, which we will indicate as ν_AMI-Ca2+_. We fit the 2DIR data with three Gaussian-shaped peaks to extract the relative area of the ν_AMI-Ca2+_ amide band at different concentrations of calcium ions. In this fit, we used two Gaussian-shaped peaks to describe the ν_ant_ and ν_AM.I_ vibrational bands in the absence of calcium, and a third Gaussian-shaped peak to describe ν_AMI-Ca2+_ (see Supplementary Methods for details). The central frequencies and the widths of the three bands were global parameters in the fit, meaning they were fixed at all studied calcium concentrations, and only the amplitudes of the three bands were allowed to be different at different calcium concentrations. Examples of the fits are reported in Figs. S2-3. We assume that ν_AMI-Ca2+_ and ν_AM.I_ have the same cross-section, and thus the fraction of amide groups bonded to calcium ions follows directly from the areas of the ν_AM.I_ and ν_AMI-Ca2+_ bands (Fig. 2c). We observe that at a low calcium concentration of 10 mM, a significant fraction of amide groups is already bonded to calcium ions. Due to limited experimental sensitivity, measurements at physiological calcium conditions (~1-2 mM) were not possible. The fraction of amide bonded groups rises quickly with increasing calcium concentration but effectively saturates at a fraction of ~10-15 mol % of *N*-acetylglucosamine.

The observed saturation implies that the binding between Ca^2+^ and hyaluronan is best described with a model that accounts for a high affinity for calcium ions at low concentrations, but includes an energetic penalty for further binding upon complex formation. This energy penalty depends on the fraction of formed complexes, and can be accounted for by using an expression for the association equilibrium constant *E*_p_ (normalized by the thermal energy, *k*_b_*T*) weighted by the fraction of occupied binding sites *f*:

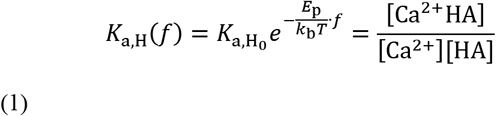

with

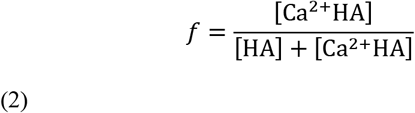

Here, [Ca^2+^] is the concentration of free calcium ions, and [HA] and [Ca^2+^HA] are the concentrations of unoccupied and occupied Ca^2+^ binding sites on HA (assuming one binding site per HA disaccharide), respectively. Using [Ca^2+^] + [Ca^2+^HA] = [Ca^2+^]_*i*_ where [Ca^2+^]_i_ is the total concentration of calcium ions in the solution and the index *i* runs over all concentrations investigated, and [HA] + [Ca^2+^HA] = [HA]_0_ where [HA]_0_ is the total concentration of binding sites, we can re-write the equilibrium expression (1) as:

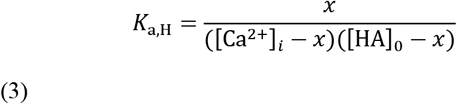

where *x* = [Ca^2+^HA]. Solving Eq. (3) for *x* and using expression (2) we obtain:

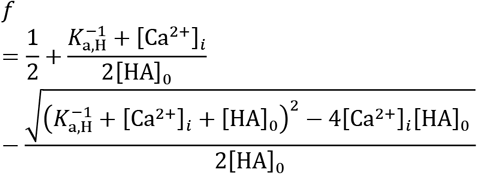

that contains an exponential term with a constant energy term (4)

To extract the zero-concentration binding constant 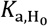, and the penalty energy *E*_p_, we globally minimize:

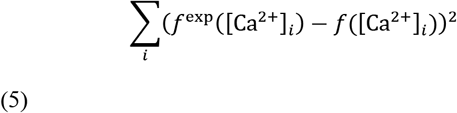

where the *f*^exp^([Ca^2+^]_*i*_) values are obtained from the 2DIR experiments. The *f*([Ca^2+^]_*i*_) values follow from solving the coupled equations (1) and (4) (see Supplementary Methods for details), where we fit the fraction of bonded amide. We obtain 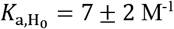, and *E*_p_ = 30 ± 5 *k*_b_*T*. The result of the fit is shown in Figure 2c.

### Effect of calcium ions on the persistence length of HA

To assess the effect of ions on the persistence length of HA at the level of individual chains, we devised an atomic force microscopy (AFM) assay to probe the stretching of individual HA chains under tensile force, as schematically illustrated in Fig. 3a (see Supplementary Methods for details). Force spectroscopy with polymer chains benefits from a uniform population of molecules with well-defined characteristics. Here we made quasi-monodisperse (*i.e.*, size distribution approaching the ideal) HA polymers with a ‘handle’ only at a single specified point. In the polysaccharide realm, our HA probe is much more homogeneous than previous naturally occurring and semi-synthetic preparations.^35^

**Figure 3:**
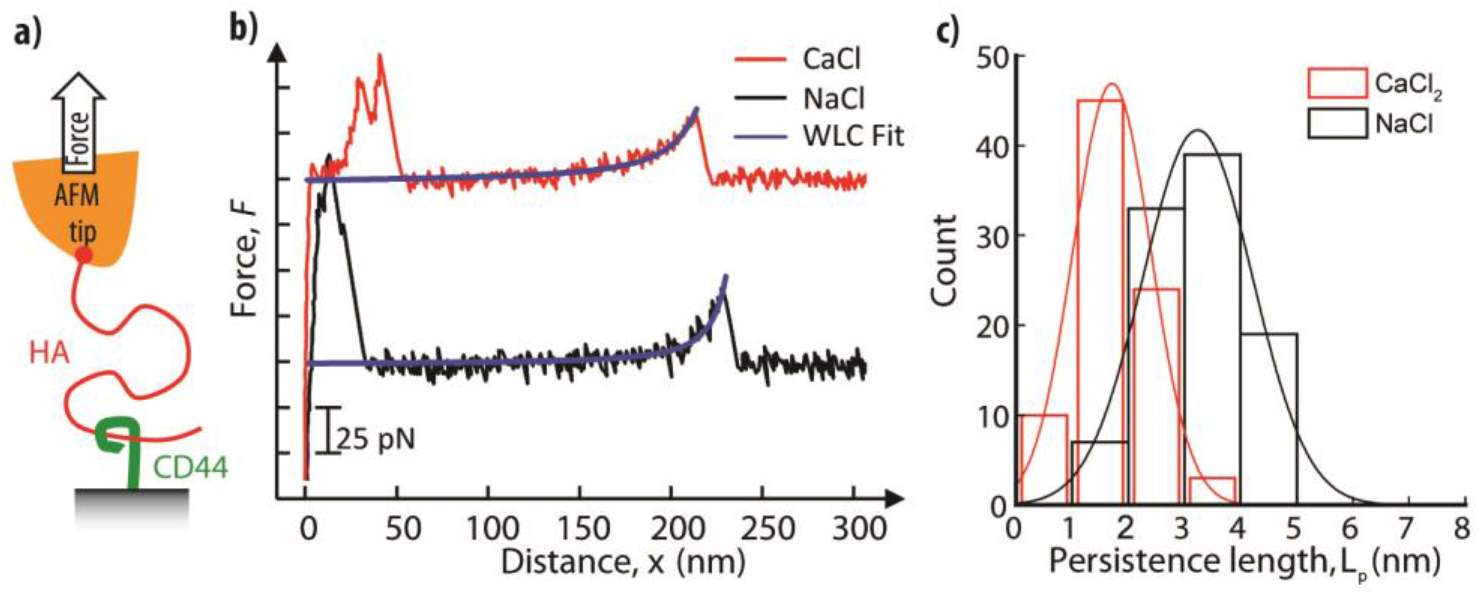
a) Illustration of the setup to measure the persistence length of HA by single molecule force spectroscopy: HA polysaccharide chains (red; *M*_W_ = 647 kDa) were grafted *via* their non-reducing end to an AFM tip; extracellular domains of the HA receptor CD44 were anchored *via* their C-terminal end to a planar support, and served as baits to capture HA chains. b) Representative force curves obtained for stretching a single HA chain in CaCl_2_ (red curve) and NaCl (black curve; offset by 100 pN along the *y-*axis for clarity). Except for a region of non-specific binding at small separations (< 50 nm), the data are well-fit by the worm-like chain (WLC) model (blue curves). c) Histograms of persistence lengths *L*_p_ determined from WLC model fits for CaCl_2_ (red bars) and NaCl (black bars). These are well-fit by Gaussian-shaped curves (lines in matching colors), giving mean and standard deviations of *L*_p_ = 3.2 ± 1.0 nm in NaCl and 1.7 ± 0.7 nm in CaCl_2_. Conditions: retract velocity of 1 μm/s, 150 mM NaCl or 50 mM CaCl_2_.

For the single chain force spectroscopy experiments, briefly, HA polysaccharide chains were anchored through a thiol handle (SH-HA); a sulfhydryl group was site-specifically introduced at the non-reducing end (a difficult or impossible location for all previous syntheses with HA polysaccharides due to the terminus’ relative lack of unique chemical reactivity, in contrast to the reducing end), to a gold-coated AFM probe. The AFM probe was brought into contact with a planar support displaying the HA receptor CD44, which acted as a bait to capture an HA chain dangling from the AFM tip. Pulling the AFM probe away from the planar support then generated a tensile force that was monitored as a function of the probe-support distance, as exemplified in Fig. 3b. Care was taken to adjust the surface densities of HA and CD44 such that the rupture of individual HA•CD44 bonds, and thus the stretching of individual HA•CD44 bonds, and thus the stretching of individual HA chains, could be resolved (Fig. S4). The force *vs.* extension curves could be well-fitted with the worm-like chain (WLC) model (Fig. 3b; see also Supplementary Methods for details), as expected for flexible and sufficiently long polymer chains. Histograms of the persistence lengths *L*_p_ extracted from these fits (Fig. 3c) show that calcium ions decrease the persistence length of HA. At physiological concentrations of monovalent salt (150 mM NaCl), we found *L*_p_ = 3.2 ± 1.0 nm, in reasonable agreement with previous work.^36^ In the presence of 50 mM CaCl_2_, the persistence length was reduced almost two-fold, to 1.7 ± 0.7 nm. Such a marked decrease in persistence length, or equivalently, increase in chain flexibility, indicates that calcium ions strongly affect the molecular conformation of hyaluronan.

### Effect of calcium ions on the molecular conformation of hyaluronan

To understand why calcium condensation causes an increased chain flexibility, we performed force field-based molecular dynamics (MD) simulations with atomistic resolution and an explicit description of the solvent molecules. We performed MD simulations of aqueous solvated HA at a concentration of 50 mM CaCl_2_, as used in the AFM experiments. We used specialized and previously tested force field parameters for a short HA oligomer dissolved in NaCl and CaCl_2_ solvent environments. We employed two different sets of parameters for Ca^2+^, referred to as CaCl_2_-Deublein and CaCl_2_-OPLS (see Supplementary Methods for details), since the treatment of divalent cations is known to be challenging at the classical, *i.e.*, force-field, level of theory.^32,37^ We ran 200 ns of unbiased MD simulations of HA oligomers with a length of 8 disaccharides, starting from a straight-chain state*, i.e.* without chain bending, in the presence of either calcium or sodium ions. The last 50 ns were used for analysis, in particular to probe the effect of the cations on the structure and dynamics of the hydrogen-bonds within the HA oligosaccharide.

In Fig. 4a, we show the structure of the oligomer in which we also indicate the five studied hydrogen bonds (labeled A to E). In addition, we also studied the geometry of the complex formed with the cation (labeled F). In Fig. 4b, we show 2D histograms of the donor-acceptor distances and angles of the hydrogen bonds A to E and the amide/carboxylate/cation complex in a NaCl environment. Strong hydrogen bonds are characterized by donor-acceptor distances shorter than 3 Å and angles larger than 2.5 rad^38^, the latter implying a high degree of alignment between the acceptor, the hydrogen and the donor.

**Figure 4:**
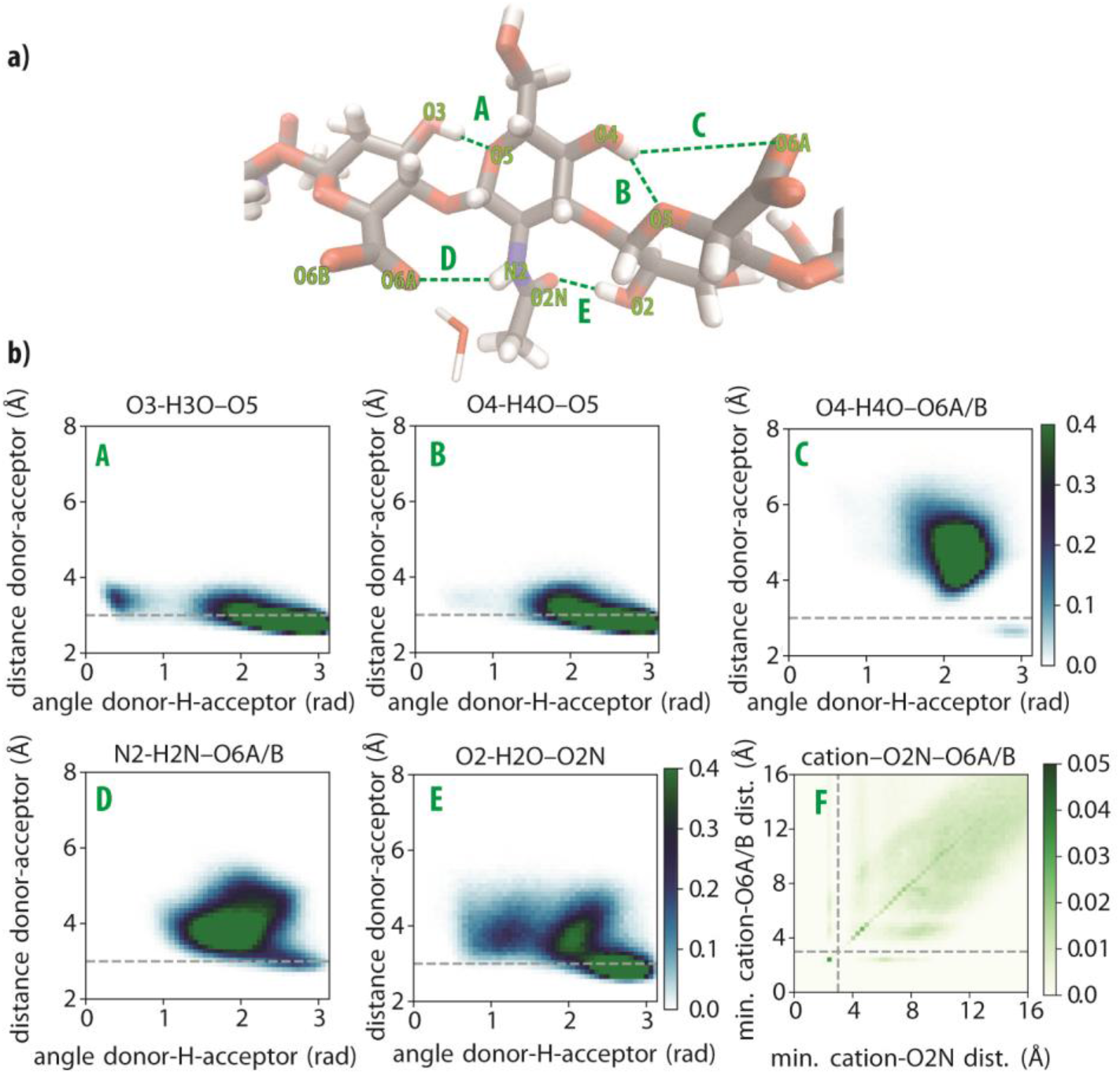
a) Key hydrogen bonds (labeled *A* to *E*) between adjacent monosaccharides along the hyaluronan chain. b) 2D histograms of the hydrogen-bond distances and angles (labeled *A* to *E*) and of the cation-amide (O2N)-carboxylate (O6A/B) complex (labeled *F*). Histograms A to E have a color bar range from 0 to 0.4 normalized units. Histogram F has a color bar range from 0 to 0.05 normalized units. The 2D histograms for the CaCl_2_-OPLS and CaCl_2_-Deublein environments are presented in Fig. S5.

In Fig. 5, we show the changes in the 2D histograms (blue for increasing, red for decreasing) when NaCl is replaced by CaCl_2_. For contacts A to E, we observe positive peaks at longer distances and smaller angles, indicating a weakening of these hydrogen bonds. An exception is hydrogen bond D, which, in addition to some shift to longer distances and smaller angles, also shows a small positive peak at a distance of 3 Å and an angle of 2.8 rad. The largest difference is observed for the complex F. The Ca^2+^ cations form close contacts with the carboxylate (<3 Å) and both close (<3 Å) and far (>3 Å) contacts with the amide. The presence of close contacts between the cation and the amide group agrees with the IR and 2DIR experimental observation that, upon addition of calcium, amide groups experience dehydration because of the formation of a direct bond between amide oxygen and the divalent ions (Figs. 1–2). The positive change at a distance of ~10 Å observed for both the amide and the carboxylate groups can be explained from cations located on neighboring monomers. The MD results thus confirm that Ca^2+^ ions have a high propensity to bind to the carboxylate and amide groups, and show that this binding results in a weakening of the intramolecular hydrogen bonds of hyaluronan.

**Figure 5:**
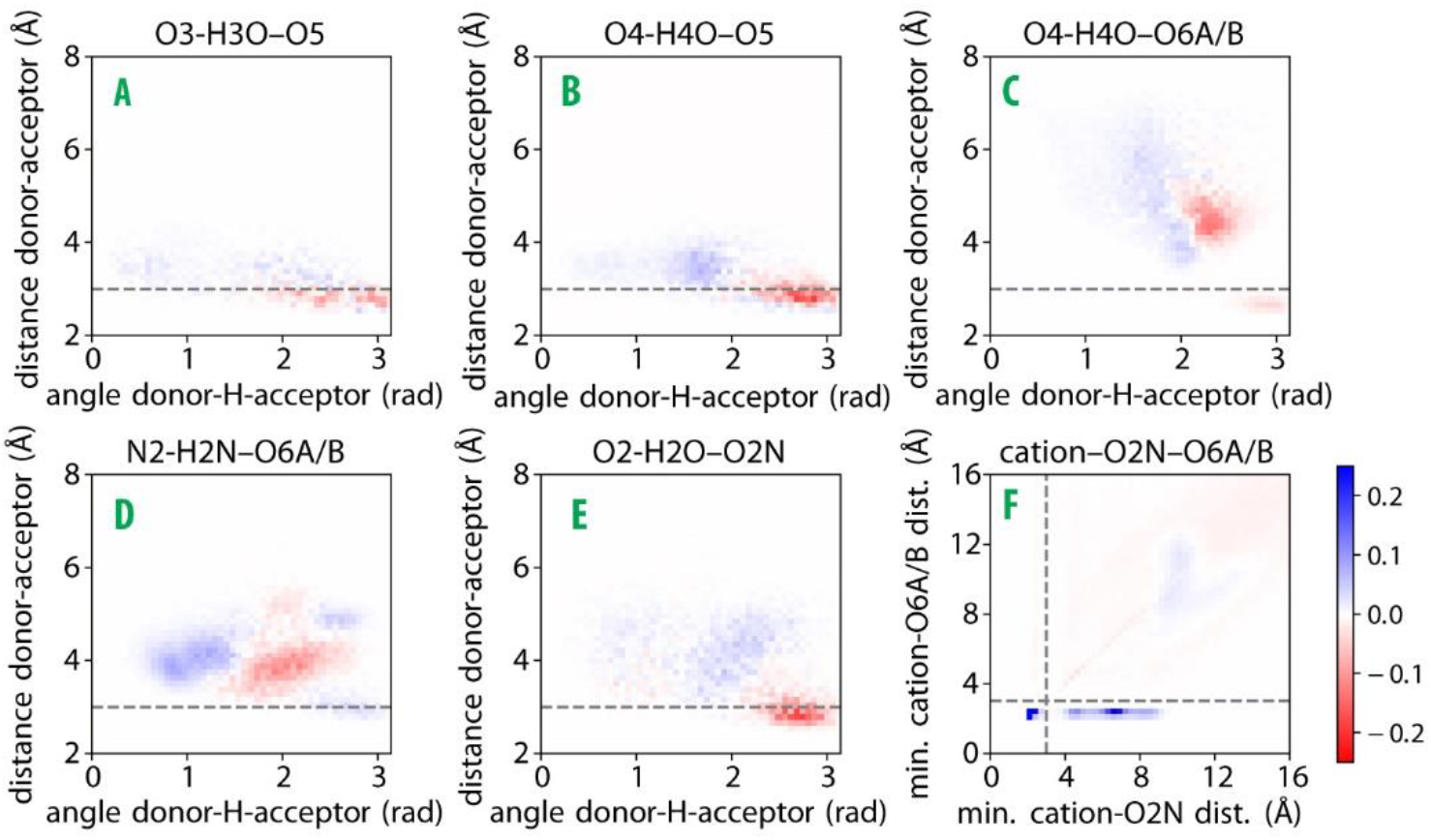
Differential 2D histograms for contacts A to F (*cf.* Fig. 4a), generated by subtracting the histograms for hyaluronan in the NaCl environment (Fig. 4b) from histograms for hyaluronan in the CaCl_2_-OPLS environment (Fig. S5a). Blue indicates an *increase* in Ca^2+^ compared to Na^+^ and red indicates a *decrease*. The differential 2D histograms obtained for the CaCl_2_-Deublein environment are presented in Fig. S6a.

Table 1 shows the probabilities of the formation of close contacts between the cation and the amide or the carboxylate group, or with both groups. For both force-fields of Ca^2+^, we find that the CaCl_2_ environment yields much greater probabilities of close contacts than the NaCl environment. The CaCl_2_-OPLS and CaCl_2_-Deublein yield similar values for the probability of close contacts with the amide (O2N; 17 and 18%, respectively), which is larger than the 5-10% extracted from the 2DIR experiments (Fig. 2d). Such discrepancy may arise because of the force field choice and/or because the cross-section of the amide vibrational band may change upon formation of a bond with the calcium ion. In Table 1 we also observe that the probabilities of close contacts between the cation and the carboxylate (O6A/B) differ between the two Ca^2+^ force fields. Previous literature reports that OPLS overestimates the Ca^2+^ carboxylate affinity^39^ and points to CaCl_2_-Deublein being more realistic. In Fig. S5, we show 2D histograms of the different hydrogen bond lengths and angles obtained with CaCl_2_-Deublein and CaCl_2_-OPLS.

**Table 1:**
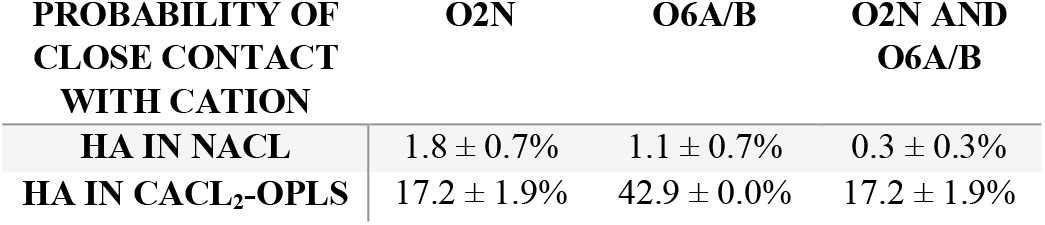
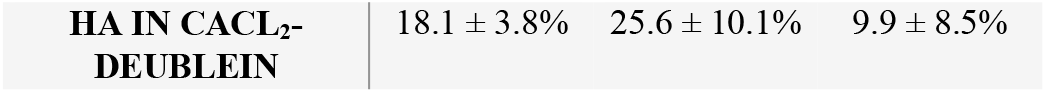
Probability (± one standard deviation) of close contact (<3 Å) of Na^+^ or Ca^2+^ cations with the carboxyl (O6A/B) or amide (O2N) oxygens, or with both oxygens simultaneously (O6A/B & O2N).

| PROBABILITY OF<br>CLOSE CONTACT<br>WITH CATION | O2N | O6A/B | O2N AND<br>O6A/B |
| --- | --- | --- | --- |
| HA IN NaCl | 1.8 ± 0.7% | 1.1 ± 0.7% | 0.3 ± 0.3% |
| HA IN CaCl<sub>2</sub>-OPLS | 17.2 ± 1.9% | 42.9 ± 0.0% | 17.2 ± 1.9% |
| **HA IN CaCl<sub>2</sub>-DEUBLEIN** | 18.1 ± 3.8% | 25.6 ± 10.1% | 9.9 ± 8.5% |

The time-averaged hyaluronan end-to-end chain length remained around 70 Å for all unbiased MD trajectories starting with a straight chain configuration (Fig. S7). This is only slightly shorter than the contour length of the HA oligosaccharide (80 Å), and suggests that large deformation events do not occur within the computational time scale and for this relatively small polymer size. In order to estimate the effect of Ca^2+^ on the flexibility of hyaluronan, we calculated the bending free energy of the same HA oligosaccharide in NaCl, CaCl_2_-Deublein and CaCl_2_-OPLS environments. This was achieved by means of a variation of the constrained MD method^40^ (see Supplementary Methods for details). The resulting bending free energy profiles, spanning end-to-end lengths from a slightly stretched (compared to the time-average length) hyaluronan oligosaccharide (75 Å) to a half-bent U-shaped one (35 Å), are shown in Fig. 6a. The two environments with Ca^2+^ are consistent with each other and present significantly lower free energies of flexed configurations than the Na^+^ environment, indicating a higher flexibility of hyaluronan in the presence of Ca^2^. For flexible polymers, the bending free energy is linearly related to the persistence length.^41^ This proportionality enables a quantitative comparison of the MD simulation results with the experimental force spectroscopy results. The free energy of the U-shaped bent HA decreases by approximately 40% from 4.96±0.73 kcal/mol in NaCl, to 3.04±0.63 kcal/mol in CaCl_2_-OPLS (or by 35% to 3.24±0.87 kcal/mol in CaCl_2_-Deublein). This reduction agrees with the approximately twofold decrease in persistence length observed by AFM (Fig. 3c).

**Figure 6:**
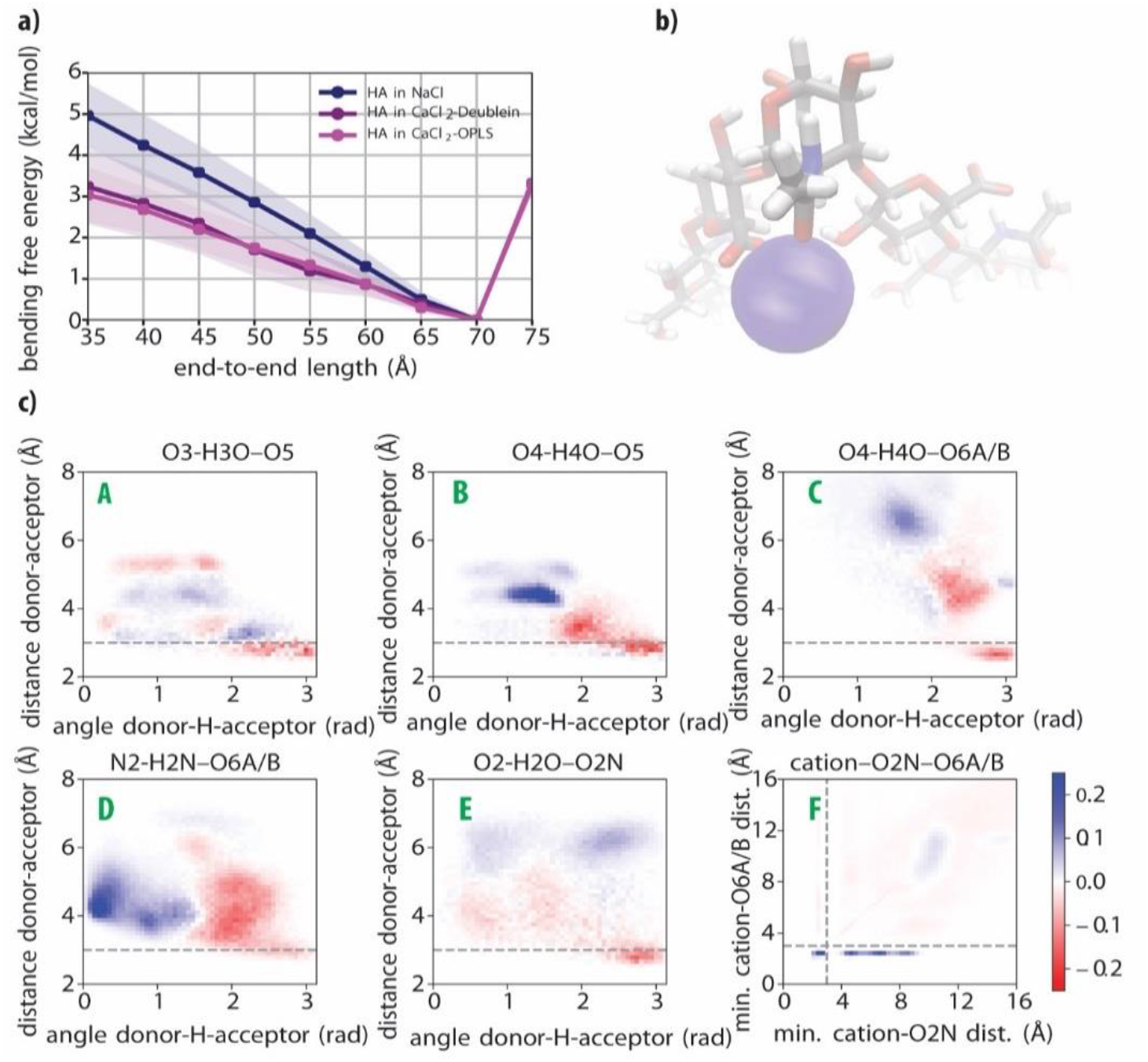
a) Bending free energy profile of hyaluronan for the three different simulation environments described in the text. The shaded regions refer to one standard deviation, as determined from a blocking analysis (see Supplementary Methods, Fig. S8 and Table S1). b) Visualization of the amide/carboxylate/cation complex rendered by averaging atomic positions during the last 25 ns of the CaCl_2_-OPLS run constrained at an end-to-end length of 35 Å. The picture is centered at the middle of the chain. The calcium is represented according to its van der Waals radius. c) Differential 2D histograms of the distances and angles in key hyaluronan inter-monosaccharide hydrogen-bonds (labeled *A to E*; *cf.* Fig. 4a) and in the cation-amide (O2N)-carboxylate (O6A/B) complex (labeled *F*) generated by subtracting histograms obtained in the NaCl simulation environment from histograms obtained in the CaCl_2_-OPLS simulation environment, with both systems constrained at an end-to-end length of 35 Å. The differential 2D histogram obtained in the CaCl_2_-Deublein environment is presented in Fig. S6b.

In Fig. 6c, the differences in the five key hydrogen bonds are analyzed for the bent configuration with an end-to-end length of 35 Å for both CaCl_2_-OPLS and NaCl. There is a general weakening of all contacts, even more considerable than the weakening observed for the unconstrained chain in Fig. 5. Contacts B to E shift to either longer distances (>4 Å) or smaller angles (<2 rad), showing a clear weakening of these hydrogen bonds. Contact A is the only hydrogen bond with a small positive change at close distances. The bending is thus accompanied by a weakening of most intramolecular hydrogen bonds.

## DISCUSSION

The 2DIR and MD results show that the HA polymer chains bind calcium ions at millimolar concentrations, leading to the formation of specific calcium complexes with amide and carboxylate groups of adjacent saccharide units. The force spectroscopy measurement and MD simulations consistently show an increase in hyaluronan flexibility in the presence of calcium. The combination of the three techniques provides a direct link between the molecular mechanism of calcium binding and its effect on the hyaluronan chain mechanics.

The 2DIR results show that the association constant of Ca^2+^ and hyaluronan has a relatively high value of 7 M^−1^ at low Ca^2+^ concentrations. This association constant is much higher than the association constant for the binding of Ca^2+^ to amide groups, for which an association constant of ~0.1 M^−1^ has been reported.^33^ We observed a similar association constant for Ca^2+^ and *N*-acetylglucosamine (0.013 ± 0.05 M^−1^, see Supplementary Methods and Fig. S9). This difference can be explained from the fact that in case of hyaluronan, the Ca^2+^ not only binds to an amide group but at the same time to a nearby carboxylate anion group. The binding with the latter group will be rather strong, thus explaining the much larger association constant of Ca^2+^ to hyaluronan compared to isolated amide groups.

It is also interesting to compare the association constant of Ca^2+^ and hyaluronan with the second saccharide unit that constitutes the building block of hyaluronan, glucuronic acid. For glucuronic acid, we found an association constant of 1.2 ± 0.2 M^−1^ (Supplementary Methods and Fig. S10), which is similar to the association constants found for complexes of simple acids and calcium^42^. The association constant of glucuronic acid and Ca^2+^ is thus approximately six times smaller than that of hyaluronan and Ca^2^. This finding indicates that in hyaluronan the relative position and orientation of the amide group of *N*-acetylglucosamine and the carboxylate anion of glucuronic acid lead to a highly favorable binding of Ca^2+^.

The strong affinity of hyaluronan for calcium may also be due to the size-charge requirements of the calcium ions. Due to its larger diameter compared to other cations, such as magnesium, the calcium ion binds well to less polar oxygens than water oxygens. Hence, the favorable inner chelation with other less polar groups, such as the amide oxygen in this case, greatly increases the stability of the calcium complex.^43–46^ It is likely that the restrained conformational fluctuations and the size-charge requirements add up synergistically, explaining the high affinity of hyaluronan to calcium. Interestingly, we find that the high affinity of hyaluronan for Ca^2+^ rapidly drops when the concentration of Ca^2+^ increases, *i.e.*, we observe that the occupation of the binding sites saturates at an occupied fraction of ~10-15%. This strong saturation is not observed for glucuronic acid. It thus appears that the binding of a calcium ion at a particular binding site of HA hinders the binding of other Ca^2+^ ions at nearby binding sites. This hindrance may be due to a conformational change of the hyaluronan induced by calcium binding, with the result that the neighboring binding sites no longer possess the highly favorable conformation of the carboxylate and amide groups that caused the initial high association constant of 7 M^−1^. This explanation is supported by MD simulations that show that the binding of Ca^2+^ induces a weakening of the hydrogen bonds and increases the flexibility of the polymer chain. The increase of the flexibility, which is also borne out by the force measurements, implies that the conformational fluctuations increase in amplitude. As a result, the time fraction in which the conformation of the binding site is favorable for binding Ca^2+^ is reduced, thereby decreasing the association constant. An additional contribution to the decrease of the affinity for Ca^2+^ may originate from electrostatic repulsion between Ca^2+^ ions at neighboring units.

## CONCLUSION

In this work, we have studied the interaction between hyaluronan polymers and Ca^2+^ ions with a combination of linear infrared spectroscopy, two-dimensional infrared spectroscopy, molecular-scale force measurements and molecular dynamics simulations. We find that hyaluronan binds Ca^2+^ with an affinity that is unusually high for inorganic ions, with an association constant of 7 ± 2 M^−1^ in the limit of millimolar Ca^2+^ concentrations. This association constant is ~6 times higher than that of glucuronic acid, which contains the same carboxylate anion motif as hyaluronan. This finding, as well as our spectroscopic data, indicates that the relative position and orientation of the amide group and the carboxylate anion groups of hyaluronan are highly favorable for binding of Ca^2+^. The molecular dynamics simulations confirm that hyaluronan has a high affinity for Ca^2+^ ions. We also observe a strong saturation of the binding of Ca^2+^ to hyaluronan at higher Ca^2+^ concentrations. This saturation effect can be well modeled with a free-energy penalty that scales with the fraction of bound Ca^2+^. The decreased binding affinity can be explained from the increase of the flexibility of the hyaluronan polymers upon the binding of Ca^2+^, as shown by the molecular dynamics simulations. An additional contribution to the saturation effect may come from electrostatic repulsion, *i.e.*, the positive charge of Ca^2+^ repels the binding of other positive charges at nearby binding locations.

The force measurements show that the binding of Ca^2+^ leads to a large decrease of the persistence length of the hyaluronan polymers which amounts to ~50% at a calcium concentration of 50 mM. The molecular dynamics simulations explain this decrease of the persistence length in terms of the weakening of several of the intramolecular hydrogen bonds, induced by the formation of the complex of Ca^2+^ with the carboxylate and amide groups.

In summary, by using a multi-technique approach, we show that a selective and localized cation binding process takes place between calcium and hyaluronan polymers, leading to the formation of specific complexes. Here we provide a detailed molecular picture of ion condensation on a polymer highlighting the severe effect of few, selective and confined electrostatic interactions on the rigidity of a polyelectrolyte chain. As the extracellular matrix contains calcium, this ion’s effect on the structure of HA chains and thus their binding to hyaladherin proteins and receptors should be considered. Moreover, given the vast employment of glycosaminoglycans to devise hydrogels with tailored applications, such as drug delivery, our findings may lead to novel ideas for creating smart materials by exploiting the unique structural properties that can be tuned by the addition of specific ions.

## ASSOCIATED CONTENT

Supplementary Methods with details on: linear and non-linear infrared spectroscopy experiments and data analysis; single-HA-chain stretching experiments and data analysis; MD simulations and their validation. Supplementary Figures S1-10. Supplementary References.

## AUTHOR INFORMATION

### Author Contributions

G.H.K. conceived the idea. H.J.B., G.G., A.P.A.O., R.P.R, G.H.K., and B.E. designed the project. G.G. performed the linear IR and 2DIR measurements, and together with H.J.B. analyzed the results. A.P.A.O. performed the MD simulations, and together with B.E. analyzed the results. F.B. performed the single chain force spectroscopy experiments and together with R.P.R. analyzed the results. X.Z., R.J.L., D.E.G. and P.L.D. produced the thio-donor and the SH-HA construct for the single chain force spectroscopy experiments. H.J.B., G.G., A.P.A.O., F.B., R.P.R., G.H.K. P.L.D., and B.E. wrote the paper. All authors agree on the contents.

‡These authors contributed equally.

## ACKNOWLEDGMENT

This work is part of the industrial partnership program Hybrid Soft Materials that is carried out under an agreement between Unilever Research and the Netherlands Organisation for Scientific Research (NWO). The authors wish to thank O. O. Sofronov (AMOLF Institute) for suggestions on the interpretation of the 2DIR and IR data, and F. Burla for useful discussions. We thank S. Banerji and D. G. Jackson (University of Oxford) for kindly providing the extracellular domain of CD44.

