## Supplementary Information for "Strong reduction of the chain rigidity of hyaluronan by selective binding of Ca^2+^ ions"

**This PDF file includes:**

Supplementary Methods
Supplementary Figures S1 to 10
Supplementary References

### SUPPLEMENTARY METHODS

#### Linear and non-linear infrared spectroscopy

***Sample preparation.*** The samples were prepared in a glass vial with heavy water (99.9% $D_2O$; Cambridge Isotope, Tewksbury, Massachusetts, USA ) and desired amounts of anhydrous $CaCl_2$ (Sigma Aldrich, St. Louis, Missouri, USA) to achieve calcium concentrations ranging from 0 to 300 mM. Hyaluronic acid produced in *Streptococcus equii*, and purified as a sodium salt in powder form ($M_w$ ~ 100 kDa; Lifecore Biomedical,Chaska, Minnesota, USA) was added to each solution to obtain a final concentration of 20 mg/ml. Samples were left to equilibrate at room temperature for 24 h before performing the measurements. Samples were then stored at a temperature of 4 °C, and used within 1 week to minimize any adverse effects of hydrolytic degradation on the experiments. The pH of the solutions was measured to be around 6.5~7, independently from the calcium concentration. The solution pH values were measured by using a pH-meter (Mettler Toledo FE20/EL20) that is calibrated for measuring the pH in $H_2O$ solutions instead of $D_2O$ solutions. The measured pH* of a $D_2O$ solution is transformed to the pH value by using the following equation: pH= (pH* + 0.4) × 0.929.

***Fourier transform infrared (FTIR) and Attenuated Total Reflection (ATR)-FTIR spectroscopy.*** All absorption measurements were performed using a Bruker Vertex 80v FTIR spectrometer equipped with a liquid-nitrogen-cooled-mercury-cadmium-telluride (MCT) detector. In all the transmission mode measurements, a standard sample cell with path length of 100 µm was used. Measurements were recorded in reflection mode by employing a commercial ATR unit (Platinum ATR Diamond). The optical lengths probed in the ATR geometry is determined by the decay length of the evanescent field, which is a function of wavelength, incident angle and the refractive indexes of the ATR crystal and the sample. At around 1600 $cm^{-1}$, the probed length is around 1 µm. In both transmission and reflection measurement modes, for every spectrum 100 scans were averaged. The spectra were recorded under nitrogen atmosphere with a wavelength resolution of 3 $cm^{-1}$. The spectra were corrected for the absorption of the solvent background at the same $CaCl_2$ concentration.

***Two-dimensional infrared (2DIR) spectroscopy.*** In the 2DIR experiments, the excitation and probe pulses were both centered at 1640 $cm^{-1}$. The experiments were carried out with a home-built setup that has been described in detail before.[1] Briefly, the excitation is performed with a pair of femtosecond mid-infrared pulses. This excitation pulse-pair induces transient absorption changes, which are monitored by a probe pulse that is delayed by a time $T_w$. After transmission through the sample, the probe pulse is sent into an infrared spectrograph and detected with an infrared mercury-cadmium-telluride (MCT) detector array, yielding the transient absorption signal as a function of the probe frequency. The dependence of the transient absorption signal on the excitation frequency is determined by measuring transient absorption signals for many different delay times between the two excitation pulses. By Fourier transformation of these spectra, the dependence of the transient absorption signal on the excitation frequency is obtained. By plotting the transient absorption signal as a function of the excitation and the probing frequency, we obtained a two-dimensional infrared (2DIR) transient absorption spectrum for each delay time $T_w$. The 2DIR spectra shown in Fig. 2 and Fig. S1 were measured at a short waiting time of 0.3 ps, because at this waiting time spectral diffusion effects are negligible.

#### Fit of the 2DIR diagonal slices

In order to extract the fraction of amide groups bonded to $Ca^{2+}$ ions, we took the isotropic 2DIR signal along the diagonal in the 2DIR spectra as a function of the probe frequency at a waiting time of 0.3 ps. All 2DIR spectra were reproduced at least three times at each $Ca^{2+}$ concentration. Fig. S1 reports representative 2DIR spectra and signals for one data set.

We fit the l diagonal slices of the 2DIR spectra using a global fitting procedure based on the minimization of the following sum of square errors:

$$E = \sum_i \left(S^{\text{fit}}([\text{Ca}^{2+}]_i) - S^{\text{exp}}([\text{Ca}^{2+}]_i)\right)^2, \tag{S1}$$

where $[\text{Ca}^{2+}]_i$ is the total concentration of calcium ions with the index $i$ running over all concentrations tested (including 0 mM) and all data sets measured (*i.e.* at least three for each $Ca^{2+}$ concentration). We define $S^{\text{fit}}$ as the sum of three Gaussian-shaped bands describing the anti-symmetric stretching vibration of the carboxylate anion group ($g^{\text{COO}^-}$), the amide I vibration of the unbonded amide group ($g_{\text{nb}}^{\text{AM.I}}$), and the amide I vibration of an amide group bonded to a calcium ion ($g_{\text{b}}^{\text{AM.I}}$):

$$\forall_i, \quad S^{\text{fit}}([\text{Ca}^{2+}]_i) = \\ = c^{\text{COO}^-}([\text{Ca}^{2+}]_i) g^{\text{COO}^-}(\omega_0^{\text{COO}^-}, \Gamma^{\text{COO}^-}) + c_{\text{nb}}^{\text{AM.I}}([\text{Ca}^{2+}]_i) g_{\text{nb}}^{\text{AM.I}}(\omega_{0,\text{nb}}^{\text{AM.I}}, \Gamma_{\text{nb}}^{\text{AM.I}}) \\ + c_{\text{b}}^{\text{AM.I}}([\text{Ca}^{2+}]_i) g_{\text{b}}^{\text{AM.I}}(\omega_{0,\text{b}}^{\text{AM.I}}, \Gamma_{\text{b}}^{\text{AM.I}}) \tag{S2}$$

where $c^{\text{COO}^-}$, $c_{\text{nb}}^{\text{AM.I}}$ and $c_{\text{b}}^{\text{AM.I}}$ are the amplitudes of the three Gaussians that depend on the calcium concentration. $\omega_0^{\text{COO}^-}$ and $\Gamma^{\text{COO}^-}$ represent the center frequency and the width of the anti-symmetric stretching vibration of the carboxylate anion group, respectively. $\omega_{0,\text{nb}}^{AM.I}$ and $\Gamma_{\text{nb}}^{\text{AM.I}}$ represent the center frequency and the width of the unbonded amide I band, respectively. $\omega_{0,\text{b}}^{\text{AM.I}}$ and $\Gamma_{\text{b}}^{\text{AM.I}}$ represent the center frequency and the width of the bonded amide I band, respectively. To reduce the number of free parameters in the fit, we constrained the center frequencies and the widths of the $g^{\text{COO}^-}$ and $g_{\text{nb}}^{\text{AM.I}}$ bands to within ± 3 $cm^{-1}$ of the values obtained by fitting the 2DIR signal with no added calcium ions (shown in Fig. S2). The center frequency and width of the $g_{\text{b}}^{\text{AM.I}}$ band were not constrained. It is clear from Eq. (S2) that only the amplitudes $c$ of the bands were allowed to change as a function of calcium concentration, whereas all $g$, $\omega_0$ and $\Gamma$ parameters were assumed to be independent of calcium concentration. All $c$, $g$, $\omega_0$ and $\Gamma$ parameters (including $c_{\text{b}}^{\text{AM.I}}(0\ \text{mM})$) were obtained with a global fit across of all datasets. In Fig. S3 we show the results of the fit for a range of selected calcium concentrations.

From the amplitudes and widths of the $g_{\text{nb}}^{\text{AM.I,}}$ and $g_{\text{b}}^{\text{AM.I}}$ bands, and assuming that the amide I vibration of an amide group bonded to calcium has the same cross-section as the amide I vibration of a non-bonded amide group, we determined the fraction of amide groups that are bonded to a calcium ion with the following expression:

$$\forall_i, \ f^{\text{exp}}([\text{Ca}^{2+}]_i) = \frac{\Gamma_{\text{b}}^{\text{AM.I}} c_{\text{b}}^{\text{AM.I}}([\text{Ca}^{2+}]_{\text{i}})}{\Gamma_{\text{b}}^{\text{AM.I}} c_{\text{b}}^{\text{AM.I}}([\text{Ca}^{2+}]_i) + \Gamma_{\text{nb}}^{\text{AM.I}} c_{\text{nb}}^{\text{AM.I}}([\text{Ca}^{2+}]_i)} \tag{S3}$$

### Model for the binding of calcium ions to *N*-acetyl-glucosamine and glucuronic acid

We describe the binding of calcium to an amide or carboxylate group of *N*-acetyl-glucosamine and glucuronic acid as:

$$\text{Ca}^{2+} + \text{M} \xrightarrow{K_{\text{a,M}}} \text{Ca}^{2+}\text{M}, \tag{S4}$$

where M represents the amide and/or carboxylate group. The equilibrium equation of this reaction is given by:

$$K_{\text{a,M}} = \frac{[\text{Ca}^{2+}\text{M}]}{[\text{Ca}^{2+}][\text{M}]}. \tag{S5}$$

By using

$$[\text{Ca}^{2+}] + [\text{Ca}^{2+}\text{M}] = [\text{Ca}^{2+}]_i, \tag{S6}$$

where $[\text{Ca}^{2+}]_i$ is the total concentration of calcium ions, and

$$[\text{M}] + [\text{Ca}^{2+}\text{M}] = [\text{M}]_0, \tag{S7}$$

where $[\text{M}]_0$ is the total concentration of molecular groups to which $Ca^{2+}$ can bind, we can write Eq. S5 as:

$$K_{\text{a,M}} = \frac{x}{([\text{Ca}^{2+}]_i - x)([\text{M}]_0 - x)}, \tag{S8}$$

where $x = [Ca^{2+}M]$.

Solving Eq. S8 for $x$ yields as the only physically meaningful solution

$$x = \frac{K_{a,M}^{-1}+[Ca^{2+}]_i+[M]_0}{2} - \frac{\sqrt{\left(K_{a,M}^{-1}+[Ca^{2+}]_i+[M]_0\right)^2-4[Ca^{2+}]_i[M]_0}}{2}. \quad (S9)$$

To extract the binding constant $K_{a,M}$, we globally minimize the chi-square

$$\sum_i \left(x([Ca^{2+}]_i)/[M]_0 - f^{\exp}([Ca^{2+}]_i)\right)^2 \quad (S10)$$

where $f^{\exp}$ is the experimentally determined fraction of bonded amide or carboxylate as obtained from Eq. S3. Experimental linear IR spectra, which were measured at least two times, and fits are reported in Fig. S7 for N-acetyl glucosamine and in Fig. S8 for glucuronic acid.

### Saturation model for the binding of calcium ions to hyaluronan

We describe the binding of calcium to hyaluronan with a binding constant $K_{a,H}$:

$$Ca^{2+} + HA \xrightarrow{K_{a,H}} Ca^{2+}HA \quad (S11)$$

The fraction of binding sites on hyaluronan that are occupied with calcium rapidly saturates with increasing added calcium concentration, indicating that the binding process is anti-cooperative. We model this effect with a penalty energy for $Ca^{2+}$ binding that increases with the fraction of occupied sites. To describe this, we take the equilibrium equation,

$$K_{a,H} = \frac{[Ca^{2+}HA]}{[Ca^{2+}][HA]}, \quad (S12)$$

and set the equilibrium constant $K_{a,H}$ equal to its value in the limit of zero occupancy $K_{a,H_0}$ multiplied with an exponential term containing a penalty energy $E_p$ that is proportional to the fraction of occupied sites:

$$K_{a,H}(f) = K_{a,H_0} e^{-\frac{E_p}{k_b T} \cdot f}, \quad (S13)$$

with

$$f = \frac{[Ca^{2+}HA]}{[HA]+[Ca^{2+}HA]}. \quad (S14)$$

As in the previous model for monomers, we can rewrite Eq. S12 as

$$K_{a,H} = \frac{x}{([Ca^{2+}]_i - x)([HA]_0 - x)}, \quad (S15)$$

where $x = [Ca^{2+}HA]$.

Solving Eq. (S15) yields as the only physically meaningful solution:

$$x = \frac{K_{a,H}^{-1}+[Ca^{2+}]_i+[HA]_0}{2} - \frac{\sqrt{\left(K_{a,H}^{-1}+[Ca^{2+}]_i+[HA]_0\right)^2-4[Ca^{2+}]_i[HA]_0}}{2}. \quad (S16)$$

By using $[HA] + [Ca^{2+}HA] = [HA]_0$ and $f = x/[HA]_0$ we obtain

$$f = \frac{1}{2} + \frac{K_{a,H}^{-1}+[Ca^{2+}]_i}{2[HA]_0} - \frac{\sqrt{\left(K_{a,H}^{-1}+[Ca^{2+}]_i+[HA]_0\right)^2-4[Ca^{2+}]_i[HA]_0}}{2[HA]_0} \quad (S17)$$

To extract the zero-concentration limit binding constant $K_{a,H_0}$, and the penalty energy $E_p$, we globally minimize:

$$\sum_i \left(f^{\exp}([Ca^{2+}]_i) - f([Ca^{2+}]_i)\right)^2, \quad (S18)$$

where the $f^{\exp}([Ca^{2+}]_i)$ are obtained from Eq. S3, and the $f([Ca^{2+}]_i)$ follow from solving the following two coupled equations:

$$\forall\ [\mathrm{Ca}^{2+}]_i,\ \begin{cases} f - \frac{1}{2} - \frac{K_{\mathrm{a,H}}^{-1}+[\mathrm{Ca}^{2+}]_i}{2[\mathrm{HA}]_0} + \frac{\sqrt{\left(K_{\mathrm{a,H}}^{-1}+[\mathrm{Ca}^{2+}]_i+[\mathrm{HA}]_0\right)^2-4[\mathrm{Ca}^{2+}]_i[\mathrm{HA}]_0}}{2[\mathrm{HA}]_0} = 0 \\ \log K_{\mathrm{a,H}} - \log K_{\mathrm{a,H_0}} + \frac{E_{\mathrm{p}}}{k_{\mathrm{b}}T} f([\mathrm{Ca}^{2+}]_i) = 0 \end{cases} \qquad \text{(S19)}$$

### Single-HA-chain stretching experiments by atomic force microscopy

***Reagents.*** A thiol-tagged hyaluronan (SH-HA) construct was synthesized in a multi-step process to make quasi-monodisperse HA chains and then a single thiol-containing monosaccharide was added to their non-reducing termini. First, synchronized polymerization with UDP-GlcNAc and UDP-GlcA donors and a HA tetrasaccharide acceptor primer was used to create the HA polysaccharide chain.[2] This purified polymer had an average $M_W$ of 667 kDa (polydispersity 1.011) according to size exclusion chromatography multi-angle laser light scattering. Second, to ensure that all HA chains had the identical terminal sugar, a single monosaccharide addition was performed ('end-capping') with 100 equivalents of UDP-GlcA (50 mM HEPES, pH 7.2, 1 mM $MnCl_2$, 1.6 mg/ml polymer, 1 mg/ml recombinant PmHAS, overnight at 30 °C; Sigma). This GlcA-capped HA precursor (GlcA-HA) was deproteinized by *n*-butanol extraction and then isolated by strong anion exchange chromatography (SAX) using a Vivapure Mini H Q spin column (Sartorious) according to the manufacturer's general instructions. The reaction mixture was applied to the spin unit equilibrated in 20 mM HEPES, pH 7.2, washed with 3 sequential steps of 50, 100, and 300 mM NaCl, and then the bound GlcA-HA polymer was eluted with 0.5-1 M NaCl steps. These latter fractions were precipitated with 70% ethanol (final) at -20 °C overnight, centrifuged (18,000 × *g*, 30 min) to harvest the polymer pellet, washed with 70% ethanol and dried at room temperature. The purified GlcA-HA was then re-suspended in water overnight at room temperature. A single monosaccharide addition was performed with 5 equivalents of a reduced thiol-containing sugar donor, UDP-4-thio-GlcNAc (manuscript in preparation), in 50 mM HEPES, pH 7.2, 1 mM $MnCl_2$, 0.5 mM DTT, 0.7 mg/ml polymer, 2 mg/ml recombinant PmHS-B [3,4] for 2 days at 30 °C over-layered with nitrogen gas. An additional aliquot of enzyme was added to 3 mg/ml final concentration and DTT was added to 3.5 mM final concentration after the first day of incubation. This SH-HA polymer was deproteinized with *n*-butanol and isolated by SAX similar to GlcA-HA. The addition of a new thiol group on the SH-HA was confirmed by reaction with fluorescein-maleimide (Invitrogen) and gel analysis (data not shown); the reaction product with SH-HA glowed as expected, but the parental 667 kDa HA polymer did not.
The extracellular domain of human CD44 with $His_{10}$ and biotin tags at the C-terminus (CD44-b) was produced recombinantly in CHO K1 cells and purified by His-tag affinity and size exclusion chromatography, as described in detail elsewhere.[5] Oligo(ethylene glycol) (OEG) constructs were purchased from Polypure (Oslo, Norway), one made of $EG_7$ with a hydroxyl group on one end and a thiol on the other end ($(OEG)_7$-SH), and the other containing $EG_{10}$ with a biotin on one end and a thiol on the other end (b-$(OEG)_{10}$-SH).
Gold-coated AFM probes with a nominal cantilever spring constant of 6 pN/nm and a nominal tip apex diameter of 30 nm (OBL Biolevers) were purchased from Bruker AFM Probes (Camarillo, CA, USA). The real spring constant for the employed cantilever batch was determined by the thermal noise method to be 8 pN/nm.

***Functionalization of AFM probe and planar substrate.*** HA-SH was anchored to atomic force microscopy (AFM) probes exploiting the strong and specific interaction between thiol and gold. The AFM probes were exposed to UV/ozone (Bioforce Nanoscience, Ames, IA) for 30 min, then immersed in a solution of 1 µM HA-SH dissolved in 1 M $KH_2PO_4$ (pH 4) for 5 min, and subsequently washed and immersed in an aqueous solution of 1 mM $(OEG)_7$-SH for 5 min to backfill and passivate the gold surface. After rinsing with ultrapure water, the functionalized probes were immersed in working buffer. This functionalization protocol has been demonstrated to generate a low coverage of functional organic molecules.[6,7] The AFM probes were used immediately after functionalization.
CD44-b was anchored through its biotin tag, located at the C-terminus to recapitulate the orientation of the extracellular receptor domain on the cell membrane, to a planar gold surface, as reported in detail previously.[5] Briefly, the gold surface was first functionalized with a mixed monolayer of b-$(OEG)_{10}$-SH and $(OEG)_7$-SH (made from an ethanolic solution with a total thiol concentration of 1 mM and a molar ratio of 98:2), and subsequently with a monolayer of streptavidin. CD44-b was then incubated, with the concentration (0.5 µg/mL) and incubation time (30 min) tuned to generate a low coverage (0.14 pmol/cm$^2$, corresponding to a root-mean-square receptor distance of 35 nm; see Supplementary Figure S3 in ref. 5 for details).

***Single-HA-chain stretching experiments*** were performed by AFM, using a NanoWizard IV system (JPK BioAFM Business, Bruker Nano GmbH, Berlin, Germany) at ambient conditions. To facilitate robust comparative analysis, measurements in NaCl and $CaCl_2$ solutions were performed sequentially using the same HA-coated AFM probe and CD44-coated substrate. The first solution contained 150 mM NaCl along with 10 mM HEPES (pH 7.4) in ultrapure water, and the second solution contained 50 mM $CaCl_2$ in ultrapure water. Care was taken to keep the AFM probe and planar substrate wet at all times throughout the experiment. Force curves were acquired at a set approach and retract velocity of $v$ = 1 μm/s, a maximal applied load of 600 pN, and a minimal surface dwell time. 500 force curves were acquired for each experimental setting.

***Data analysis.*** The force curves were analyzed with JPK Data Processing software. The persistence length of HA was quantified from single-HA-chain stretching curves, through fits with the worm-like chain (WLC) model, *i.e.*, $F(x) = k_\mathrm{b}T/L_\mathrm{p} \times [1/4(1-x/L_\mathrm{c})^2 + x/L_\mathrm{c} - 1/4\,]$,[8] with $F$ the stretching force, $x$ the distance, $L_\mathrm{p}$ the persistence length, and $L_\mathrm{c}$ the effective contour length representing the HA contour length from the anchor point on the AFM tip to the site of CD44 attachment. Only rupture events occurring at distances beyond 100 nm were considered as specific rupture events and considered for further analysis. The majority of force curves showed either no or a single specific rupture event. We sometimes observed more than one peak, and in these instances only the last peak was fit with the WLC model. Only curves that were well fit by the WLC model were used to extract the persistence length of HA. Instantaneous loading rates $r$ were computed from the effective spring constant $k_\mathrm{eff}$, corresponding to the slope of the WLC best-fit curve close to bond rupture, and the retract velocity $v$ as $r = k_\mathrm{eff}\, v$.

### Molecular Dynamics (MD) simulations

***Simulation setup and unbiased simulations.*** The system preparation was done with GROMACS 5.1.4.[9] The starting topology of hyaluronan with 8 disaccharides (where *N*-acetylglucosamine was the monosaccharide at the reducing end) was taken from the supporting data of reference.[10] The interatomic interactions of the chain were modeled with the GLYCAM06h forcefield,[11] specifically designed for polysaccharides. The oligosaccharide was solvated in a 10 × 10 × 10 $nm^3$ cubic box containing 1000 g/l water molecules, which were modeled by the SPC/E forcefield.[12] The force field parameters of water (SPC/E) were chosen for their compatibility and have been used successfully before in systems with $Ca^{2+}$ ions.[13,14]

We prepared two different simulation systems. In the first, random water molecules were replaced with $Na^+$ and $Cl^-$ until a concentration of 50 mM NaCl was reached. The $Na^+$ and $Cl^-$ atoms were treated with the AMBER99SB-ILDN force field,[15] which is widely accepted in force-field MD. In the second system, random water molecules were replaced with $Ca^{2+}$ and $Cl^-$ until a concentration of 50 mM $CaCl_2$ was reached. The divalent calcium ions are known to be difficult to parameterize with force fields.[13] We therefore used two different models for parameterization. The parameters established by Deublein and coworkers[13] (here called Deublein) were purposely designed for cations in aqueous solutions, while the general purpose OPLS-AA force field[16] (here called OPLS) has been reported to show a good match of the amide-$Ca^{2+}$ dissociation free energy with the free energy obtained with quantum mechanical density functional theory (DFT) based MD.[14] Both $Ca^{2+}$ force fields have been used before in combination with the SPC/E water model, whose lack of polarizability effects does not seem to affect the quality of the simulated structures and free energy profiles. All systems were simulated in the NPT (*i.e.* constant number of particles, pressure and temperature) ensemble with the canonical sampling through velocity rescaling (CSVR) thermostat[17], which was set to 310 K with a time constant of 0.1 ps, and the Parrinello-Rahman barostat[18], which was set to 1.0 bar with a time constant of 1.0 ps. Unbiased MD simulations were run for 200 ns, and data for analysis were taken from the last 50 ns of each run. The MD time step size was 2.0 fs. All systems were energy minimized before starting equilibration and production runs.

***Constrained simulations and free energy calculations.*** We employed a variation of the constrained MD method with stiff restraints instead of holonomic, or absolute, constraints. In this scheme, several simulations are performed, each of them with a stiff harmonic restraint centered on a particular value along a chosen order parameter, $z(\vec{q})$, which is a function of the system's atomic coordinates. If the restraints are stiff enough, and the oscillations around the centers of the restraints are negligible, after sufficient sampling the average force exerted by each restraint is an estimate of the negative of the underlying free energy

gradient at that $z$ value, $\langle F(z) \rangle \approx -\partial A/\partial z$. Subsequently, we used numerical integration to recover a free energy profile along $z$. We choose the end-to-end length as order parameter and set the stiffness restraints (with force constants of $k$ = 1000 kcal mol$^{-1}$ Å$^{-2}$ = 616 $k_b T$ Å$^{-2}$) every 5.0 Å for a range of 75 Å to 35 Å, which spans configurations from an almost fully stretched polymer, to a half-bent (U-shaped) chain.
The restraints were set using PLUMED 2.3.0.[19] We used the biasing technique of steered MD[20] to prepare the initial configurations for the constrained MD free energy calculations. We used a harmonic upper and lower wall with a force constant of $k$ = 1000 kcal mol$^{-1}$ Å$^{-2}$ = 616 $k_B T$ Å$^{-2}$ and steered it in 200 ps to the desired value for the free energy calculations. We allowed for 5 ns of equilibration to allow the conformation to relax before sampling the forces that we use for the free energy estimation. The total sampling time was 50 ns.
For the free energy estimation, the last 45 ns of sampling were divided into 9 blocks of 5 ns each, to obtain 9 free energy profiles. Fig. 6a shows the average of these profiles (solid line) and one standard deviation above and below. The increased standard deviation in the direction of the shorter end-to-end lengths is due to the fact that the integration starts from the longest distance.
To ensure that 9 blocks of 5 ns provide stable force averages, we performed a blocking analysis. We started by subdividing the last 45 ns of sampling into 90 blocks of 0.5 ns each, and then gradually increased the block size by 0.5 ns until 9 blocks of 5 ns were obtained. Fig. S8 shows the averages and standard deviations of the forces exerted by the stiff restraint in each simulation. The average force, which is the estimator of the free energy gradient, is stable independent of the block size. The standard deviation decreases significantly and appears to be stabilized at a block size of 5 ns for most cases. The only samples in which the standard deviation does not decrease are the stretched configurations with a 75 Å end-to-end length, however, these do not impact the bending free energy result as they correspond to the stretched, not the bent, chain.
To further test the free energy calculations, we also extended the simulation time for the runs constrained at end-to-end length of 35 Å in $CaCl_2$. After performing an additional 50 ns of MD, the average force exerted by the constraint did not change beyond the error bars shown in Fig. S8, confirming that the sampling time is sufficient to robustly estimate the free energy gradient.

**Table S1:**

| **Force exerted by the constraint (kcal mol $^{-1}$ Å$^{-2}$)** | **Initial 50 ns** | **Additional 50 ns** |
|---|---|---|
| **HA in $CaCl_2$-OPLS** | 0.05±0.05 | 0.05±0.06 |
| **HA in $CaCl_2$-Deublein** | 0.09±0.03 | 0.09±0.05 |

### SUPPLEMENTARY FIGURES

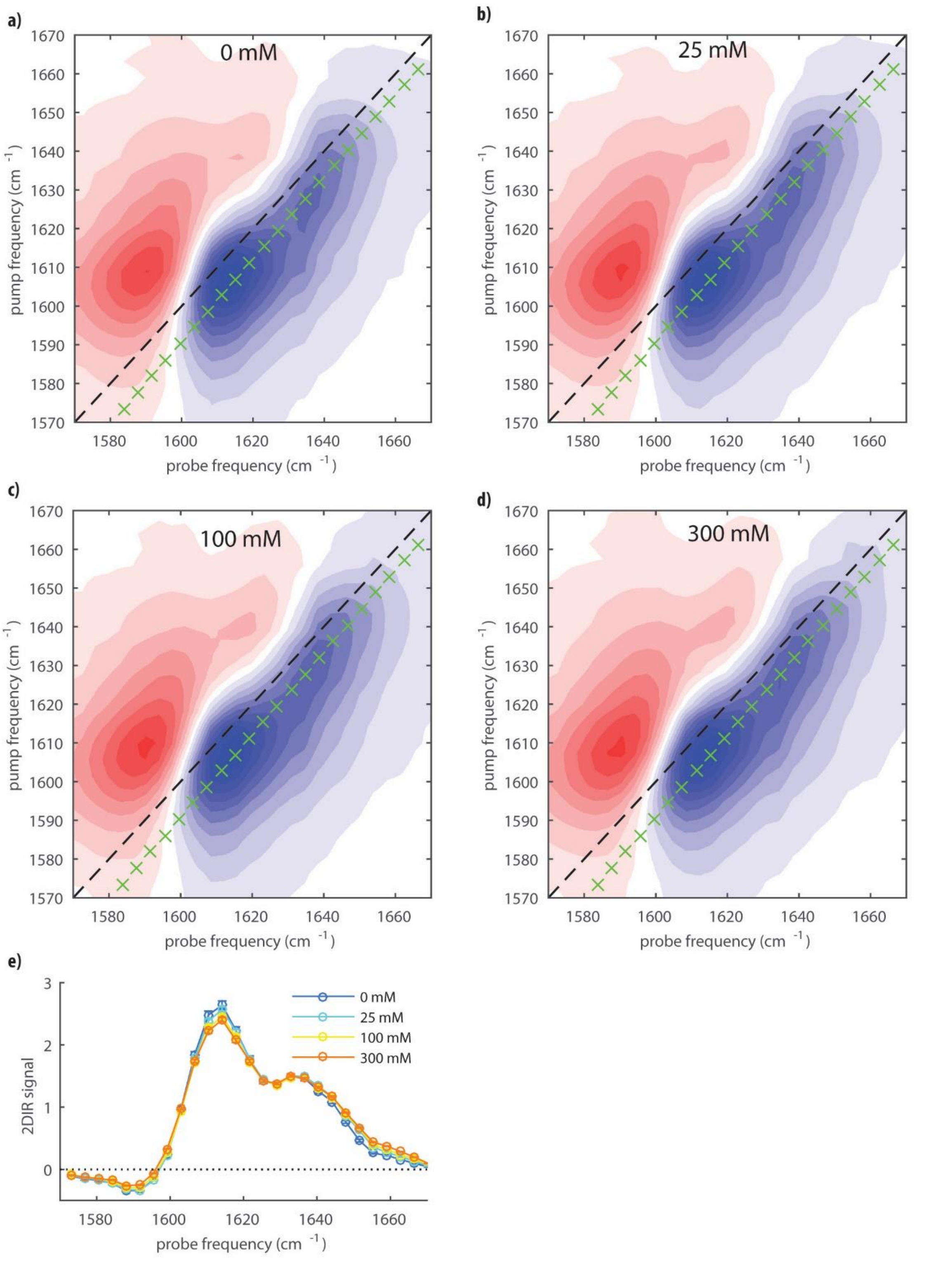


**Figure S1.** (a-d) Isotropic 2DIR spectra of hyaluronan at 20 mg/ml at different $CaCl_2$ concentrations (0 to 300 mM, as indicated in the graphs). (e) 2DIR signals taken along the diagonals marked by green symbols in a-d as a function of probe frequency.

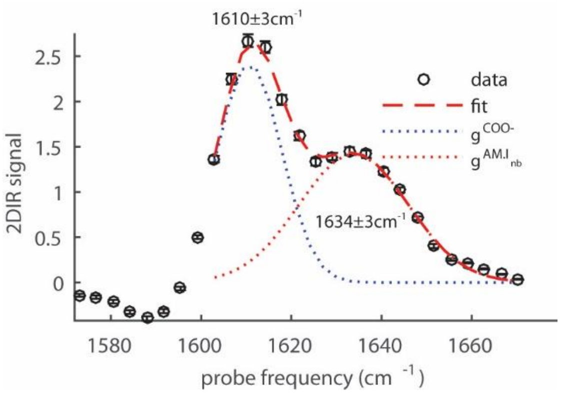


**Figure S2.** Fit of the diagonal slice of the isotropic 2DIR signal as a function of probe frequency for an aqueous solution containing 20 mg/ml hyaluronan and 0 mM $CaCl_2$. The corresponding 2DIR spectrum is shown in Fig. S1. The fit (red dashed line) to the experimental data (symbols with experimental error bars) is a superposition of two Gaussian-shaped bands describing the anti-symmetric stretching vibration of the carboxylate anion group ($\boldsymbol{g}^{\mathbf{COO^-}}$; blue dotted line), and the amide I vibration of the amide group ($\boldsymbol{g}_{\mathbf{nb}}^{\mathbf{AM.I}}$; red dotted line). We find that $\boldsymbol{\omega}_{\mathbf{0}}^{\mathbf{COO^-}} = \mathbf{1610 \pm 3\ cm^{-1}}$, $\boldsymbol{\Gamma}^{\mathbf{COO^-}} = \mathbf{24 \pm 2\ cm^{-1}}$, $\boldsymbol{\omega}_{\mathbf{0,nb}}^{\mathbf{AM.I}} = \mathbf{1634 \pm 3\ cm^{-1}}$ and $\boldsymbol{\Gamma}_{\mathbf{nb}}^{\mathbf{AM.I}} = \mathbf{40 \pm 3\ cm^{-1}}$.

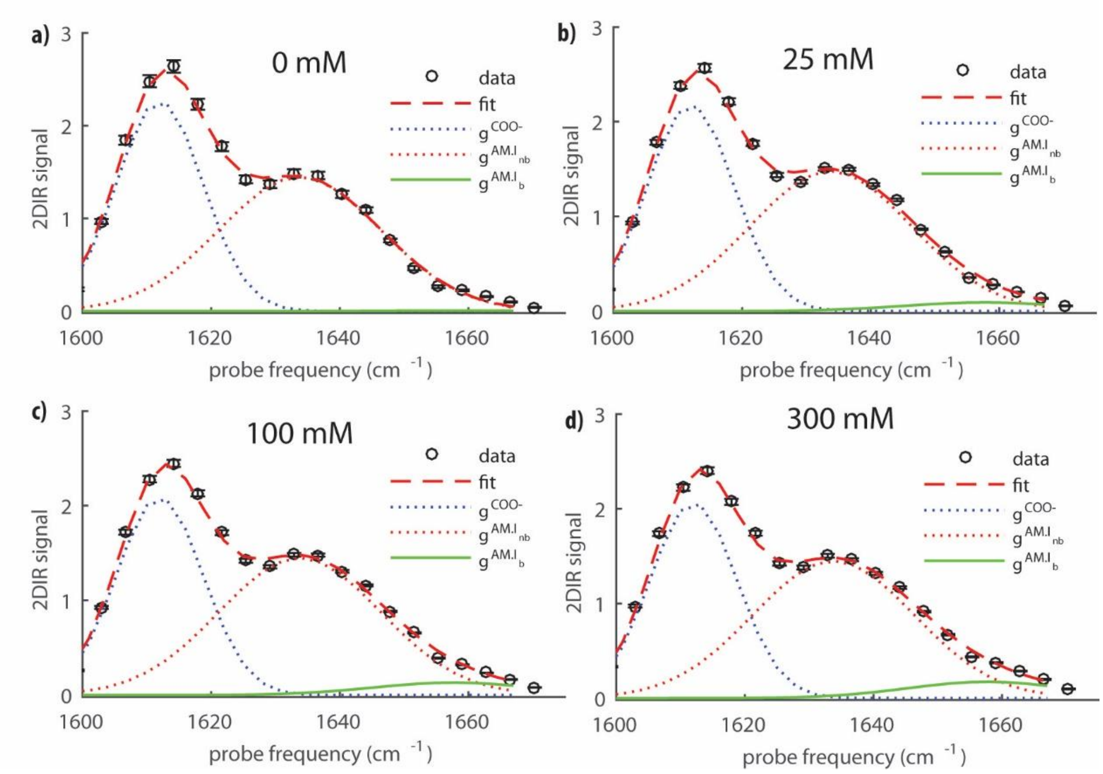


**Figure S3.** Fit of the diagonal slices of the isotropic 2DIR signals as a function of probe frequency for a solution of 20 mg/ml of hyaluronan and different $CaCl_2$ concentrations (0 to 300 mM, as indicated in the graphs). The corresponding 2DIR spectra are shown in Fig. S1. The fit is obtained by using three Gaussian-shaped bands as described in the Supplementary Methods *Fit of the 2DIR diagonal slices*. The first two Gaussians correspond to the bands shown in Fig. S2 for a solution without $CaCl_2$. The third Gaussian band represents the amide I vibrational spectrum of amide groups that are bonded to calcium. For this band we find $\boldsymbol{\omega_{0,b}^{AM.I} = 1658 \pm 3\ cm^{-1}}$ and $\boldsymbol{\Gamma_{b}^{AM.I} = 42 \pm 3\ cm^{-1}}$.

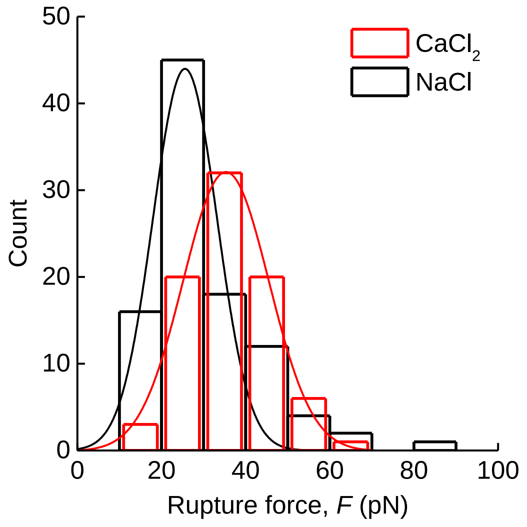


**Figure S4**. Supplementary AFM data. (a) Histograms of HA•CD44 bond rupture forces in $CaCl_2$ (red bars) and NaCl (black bars), for the data sets shown in Fig. 3. Both histograms show single peaks, and fits with Gaussian-shaped curves (lines in matching color) provide mean values and standard deviations of 35 ± 11 pN for $CaCl_2$, and 26 ± 8 pN for NaCl. The occurrence of a single peak, the comparable standard deviations between $CaCl_2$ and NaCl, and the reasonable agreement of the mean and standard deviation for NaCl with our previous work (see ref. [5]) are all consistent with single bonds breaking, and thus individual HA chains being pulled, in the majority of rupture events. Instantaneous loading rates were 1.9 ± 0.6 nN/s for $CaCl_2$ and 0.9 ± 0.4 nN/s for NaCl.

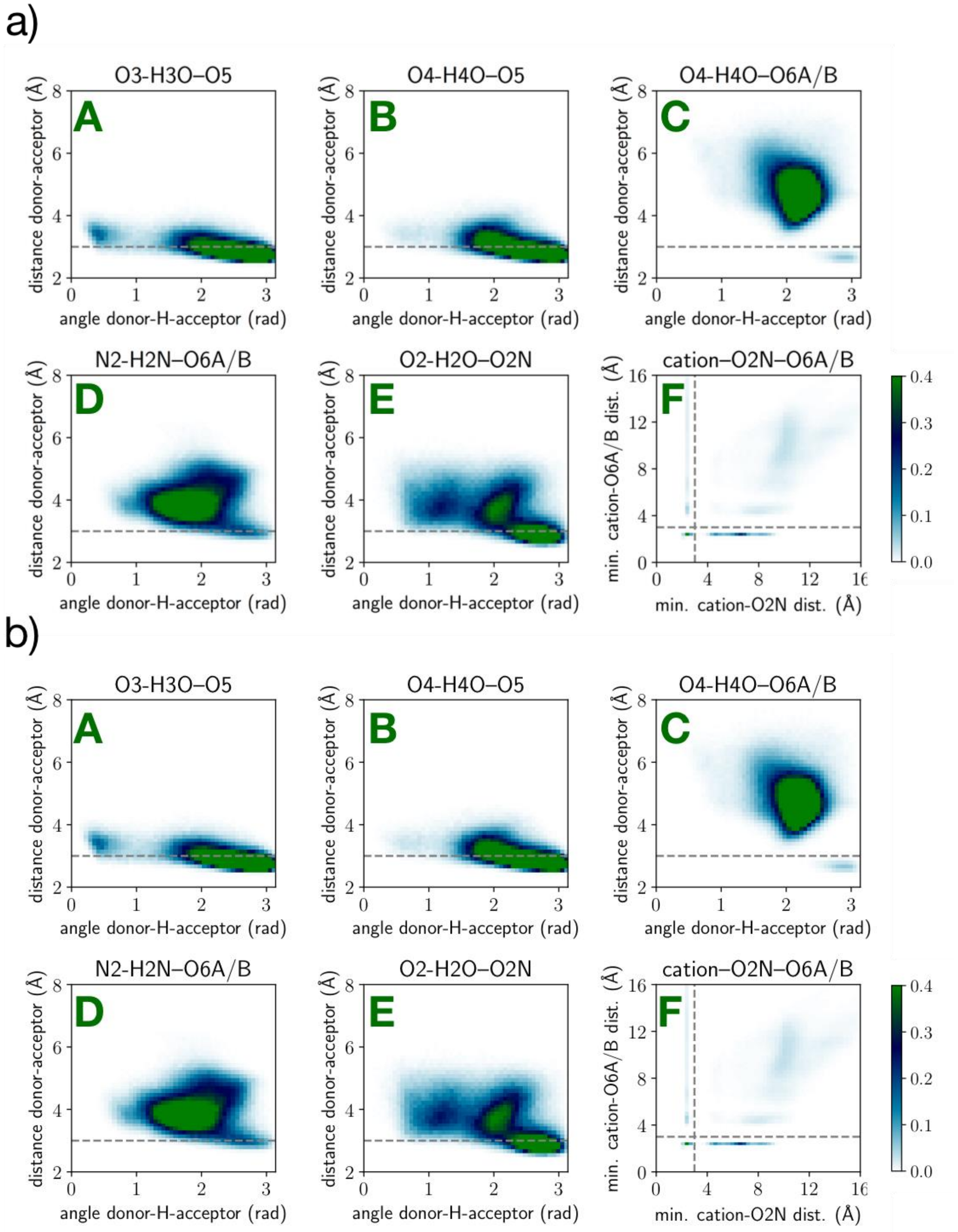


**Figure S5.** Histograms of the key hydrogen-bond distances and angles (labeled A to E) and of the cation-amide(O2N)-carboxylate(O6A/B) complex distances (labeled F) for hyaluronan in the unbiased straight-chain state in the $CaCl_2$-OPLS (a) and $CaCl_2$-Deublein (b) simulation environments. All histograms have a color bar range from 0 to 0.4 normalized units.

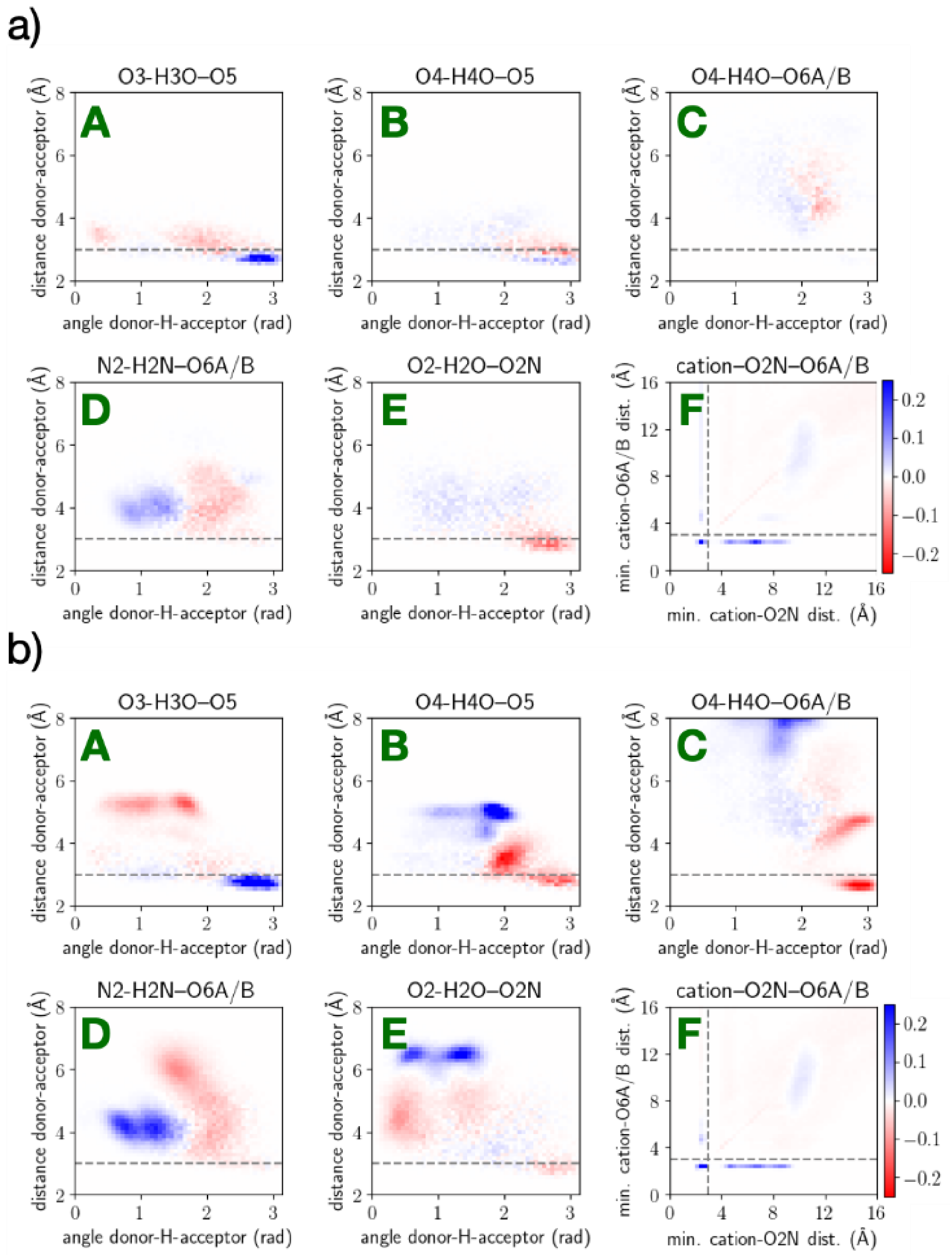


**Figure S6.** Histograms of the differences in the key hydrogen-bond distances and angles (labeled A to E) and in the cation-amide(O2N)-carboxylate(O6A/B) complex distances (labeled F) for hyaluronan in the $CaCl_2$-Deublein simulation environment compared to NaCl with both systems in the unbiased straight-chain state (a) and with both systems in the biased bent state with an end-to-end distance of 35 Å (b).

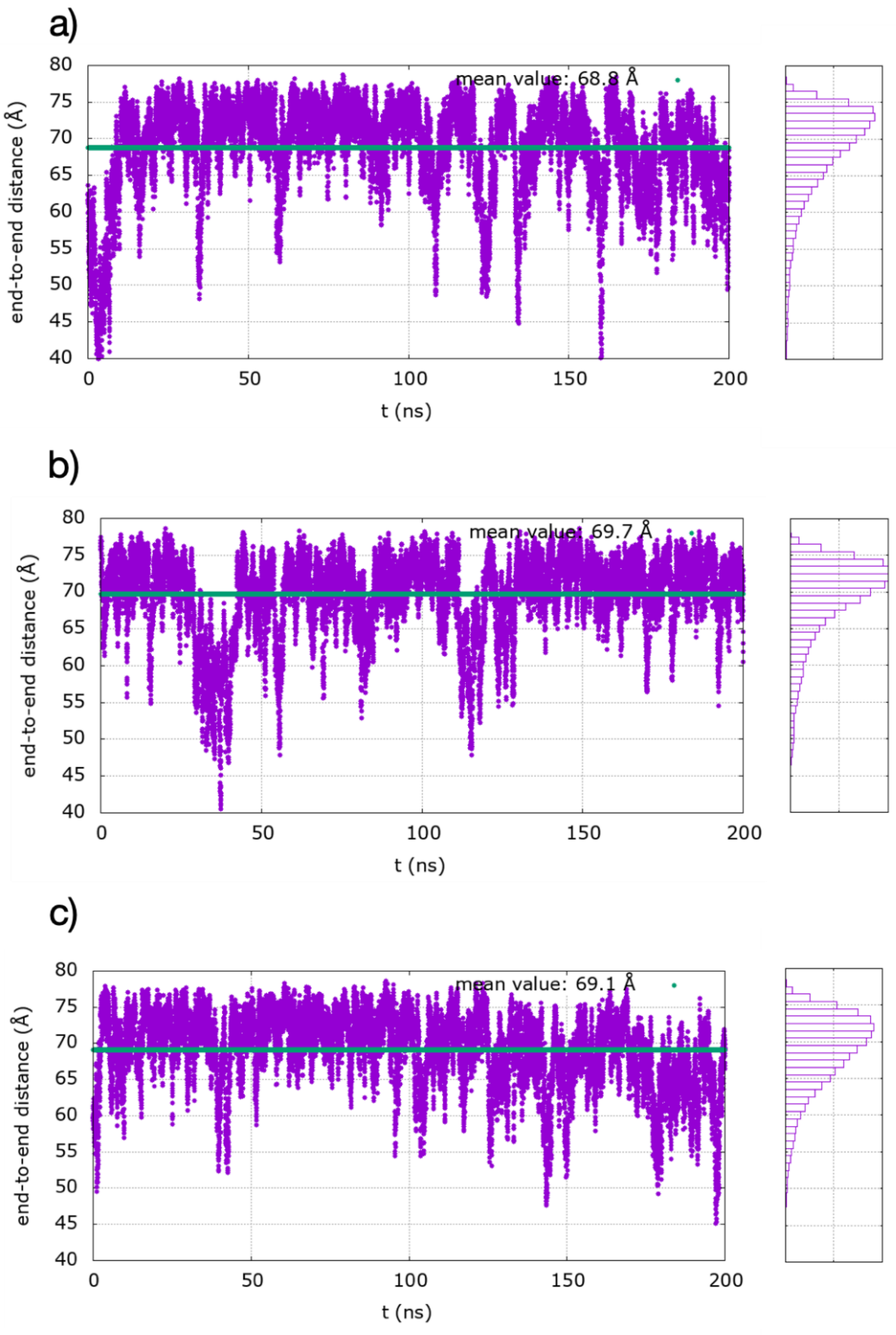


**Figure S7.** Time series of end-to-end distances (left panels), and histograms of end-to-end distances (right panels) derived from unbiased MS simulations of hyaluronan made of 8 disaccharides in the NaCl environment (a), the $CaCl_2$-OPLS environment (b), and the $CaCl_2$-Deublein environment (c). Time-averaged mean values are also indicated in the left panels (green solid lines).

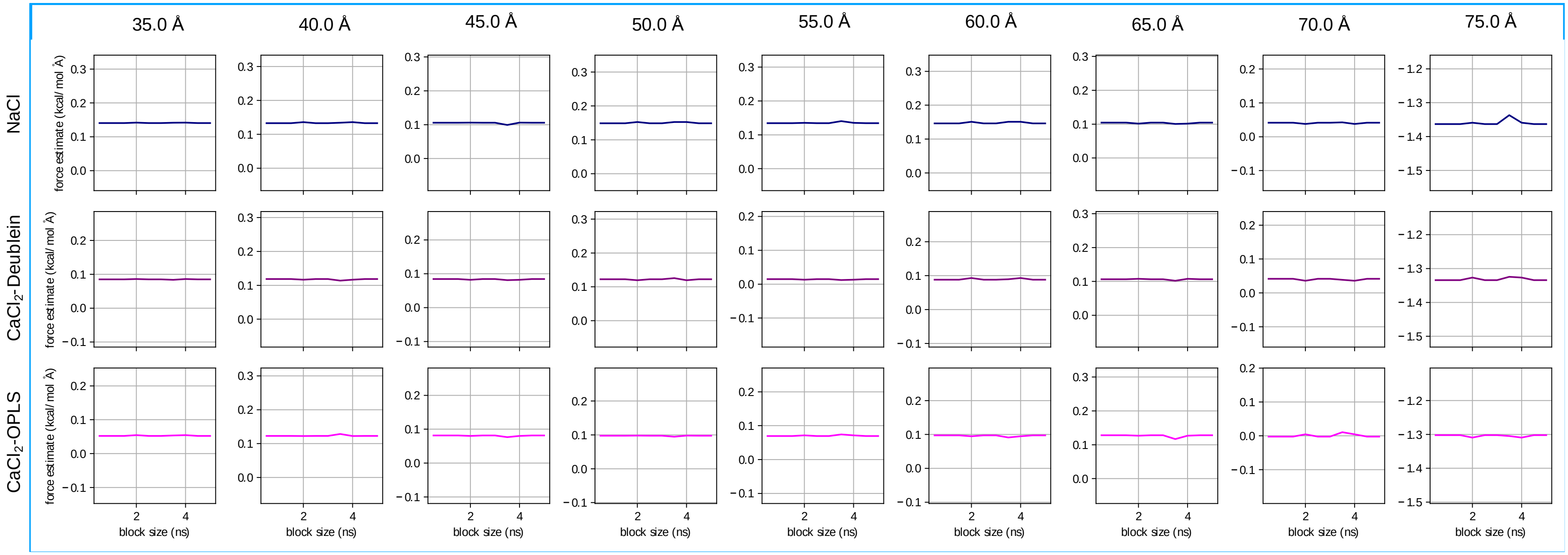


**Figure S8.** Blocking analysis of the forces exerted by each restraint (from 35 Å to 75 Å) used for the free energy calculations in the three distinct solvent environments. As a function of the block size, the solid color line shows the average force and the shaded regions show one standard deviation.

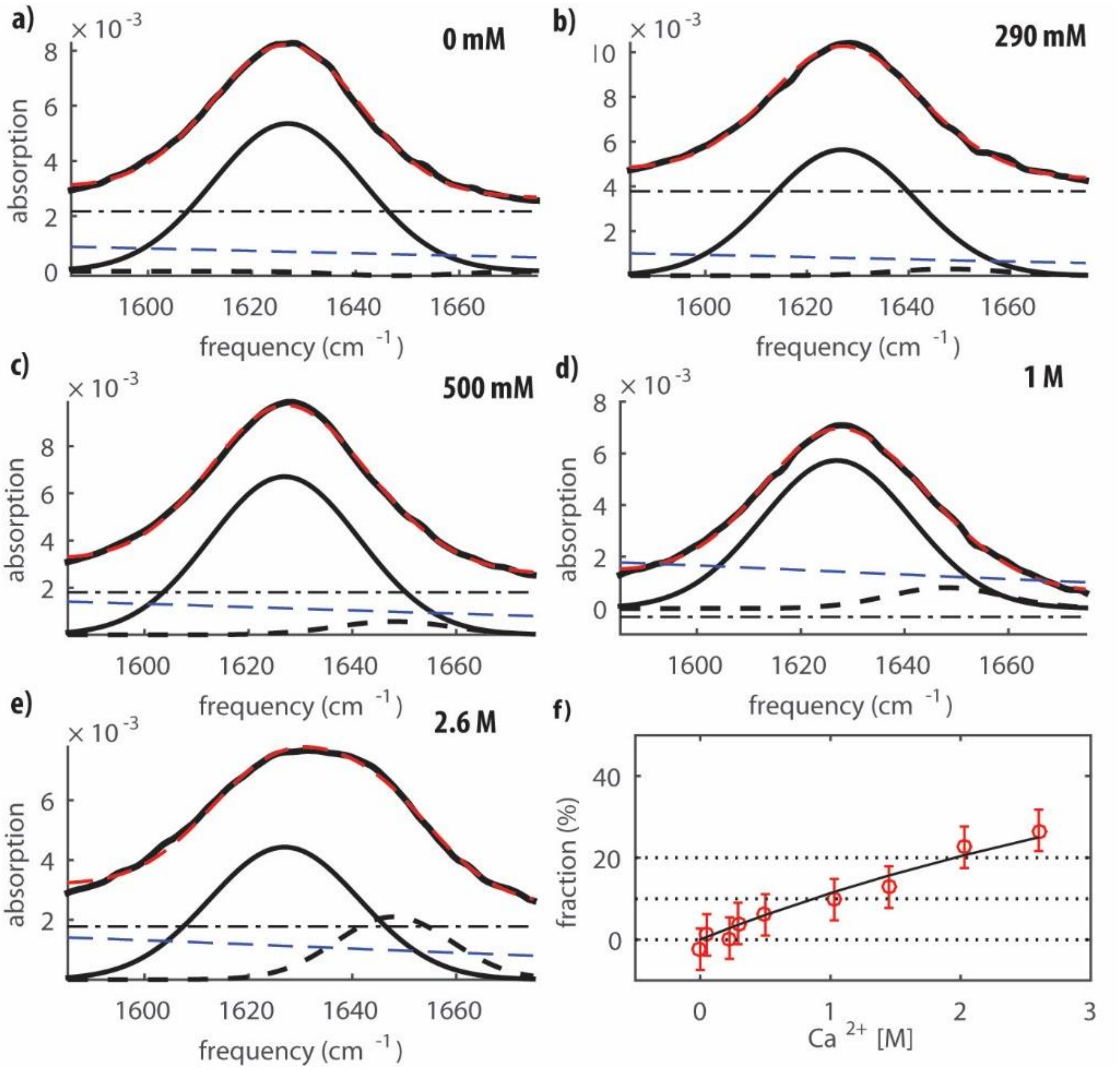


**Figure S9.** a-e) ATR-FTIR spectra (thick solid curves) of *N*-acetylglucosamine at a concentration of 10 mg/ml for six different added $CaCl_2$ concentrations (0 to 1.4 M, as indicated in the graphs). Upon addition of calcium ions, we observe a broadening and a blue shift of the band. The spectra are fitted with two Gaussian-shaped bands, which represent the amide I mode of the amide groups that are not bonded to calcium ions (solid curves) and the amide groups that are bonded to calcium ions (dashed curves). In addition to the Gaussian-shaped bands, the fit of the spectra also contains the spectrum of heavy water and a straight line (blue and black thin dashed curves, respectively) to compensate for the offset due to experimental issues encountered in the background subtraction. The Gaussian band for the non-bonded amide group is centered at 1627 ± 2 cm$^{-1}$ with a width of 37 ± 3 cm$^{-1}$, and the Gaussian band for the bonded amide group is centered at 1648 ± 3 cm$^{-1}$ with a width of 48 ± 3 cm$^{-1}$. The cumulative signals of all fit components in the global fit are also shown (thick red dashed curves) to facilitate comparison with the experimental data. In the fit, the amplitudes of the Gaussians were independently adjustable for each calcium concentration, and used to calculate the fraction of bonded amide groups. f) Fraction of bonded amide groups as a function of $CaCl_2$ concentration. The experimental fraction (red dots) is obtained by fitting the linear spectra at different concentrations, and comparing the areas of the two Gaussian-shaped bands (analogous to Eq. S3). The error-bars are obtained by propagation of the experimental errors and the fit errors. The data are fitted to the equilibrium model (Eq. S9), and we find $K_{a,M}$ = 0.13 ± 0.05 M$^{-1}$.

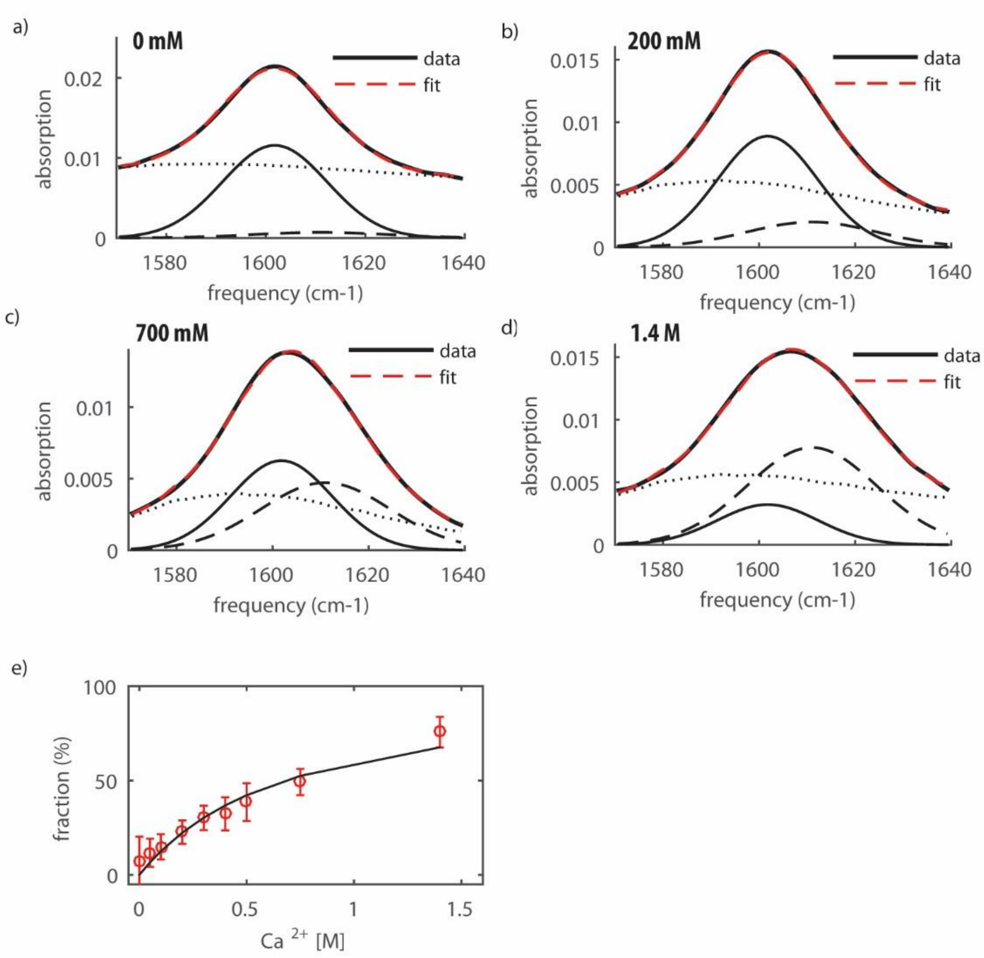


**Figure S10.** a-d) ATR-FTIR spectra of glucuronic acid (thick solid curves) at a concentration of 10 mg/ml for four different $CaCl_2$ concentrations (0 to 1.4 M, as indicated in the graphs). Upon addition of calcium ions, we observe a broadening and a blue shift of the band. The spectra are fitted with two Gaussian-shaped bands, which represent the anti-symmetric stretching modes of the carboxylate anion groups for the carboxylate groups not bonded to $Ca^{2+}$ (thin solid curves), and the carboxylate groups bonded to $Ca^{2+}$ (thin dashed curves). In addition to the Gaussian-shaped bands, the fit of the spectra also contains the spectrum of heavy water (thin dotted curves), which shows a broad band centered around 1590 $cm^{-1}$, to compensate for the offset due to experimental issues encountered in the background subtraction. The Gaussian band of the carboxylate group that is not bonded is centered at 1601 ± 2 $cm^{-1}$ and has a width of 35 ± 2 $cm^{-1}$, and the Gaussian of the carboxylate group that is bonded is centered at 1611 ± 3 $cm^{-1}$ and has a width of 45 ± 3 $cm^{-1}$. The cumulative signals of all fit components in the global fit are also shown (thick red dashed curves) to facilitate comparison with the experimental data. The amplitudes of the Gaussians are the only parameters that are allowed to vary with the added calcium concentration. e) Fraction of bonded carboxylate anions as a function of $CaCl_2$ concentration. The experimental fraction (red symbols) is obtained by fitting the linear spectra measured at different concentrations, and comparing the areas of the two Gaussian-shaped bands (analogous to Eq. S3). The error bars are obtained by propagation of the experimental errors and the fit errors. The data are fitted with the equilibrium model (Eq. S9), and we find $K_{a,M}$ = 1.2 ± 0.2 $M^{-1}$.
